# Genomic insights into the local adaptation of spontaneously occurring populations of olive trees (*Olea europaea ssp. europaea var. sylvestris*) in the Mediterranean Basin

**DOI:** 10.1101/2025.08.04.668140

**Authors:** Lison Zunino, Gautier Sarah, Lorenzo Rocchetti, Bénédicte Rhoné, Alexandre Soriano, Gaetan Droc, Pierre Mournet, Ahmed El Bakkali, Evelyne Costes, Bouchaib Khadari, Philippe Cubry

## Abstract

Living organisms are increasingly threatened by significant environmental changes, primarily driven by human-induced global alterations. Understanding and forecasting species adaptive responses to these environmental changes is therefore crucial for enhancing conservation efforts. In the Mediterranean Basin (MB), the average temperature is rising at a rate 20% faster than the global average, placing regional biodiversity at heightened risk. The study of local adaptation is thus particularly relevant. In the western MB, spontaneously occurring olive populations consist of both wild olives and admixed resulting of crop-to-wild gene flow. The wild is likely harbouring adaptive genetic variation shaped by local environmental pressures. In this study, we analysed target genomic sequencing data from spontaneously occurring populations (comprising both wild and admixed trees) as well as cultivated olive trees. We performed selective sweep and GEA analyses in wild olives to identify genomic regions associated with environmental variables. Our findings identified signatures of adaptation particularly associated with precipitation and temperature. Notably, admixed individuals retained many wild candidate SNPs in their genomes suggesting that they might retain a certain level of adaptation to their local environment. Those results are in line with recent studies suggesting hybrids could adapt more rapidly to novel environments than their parental populations.

**Key-words**: Local adaptation, selective sweep detection, crop-to-wild gene flow, *Olea europaea* L., population genomics, genome-environment association

## Introduction

Nowadays, living organisms face significant environmental changes due to human-induced global change. This phenomenon, resulting from human activities since the industrial era, has led to substantial climate shifts worldwide, particularly an increase in global average temperatures and changes in precipitation patterns. In the Mediterranean Basin (MB), global change has already been documented, with average temperatures rising 20% faster than the global average (*1*). This is noteworthy as this region is also a biodiversity hotspot, home to numerous endemic plant and animal species (*2*). Understanding how natural perennial populations respond to environmental constraints due to global changes is thus crucial. Determining the relative strength of climatic drivers that have shaped species evolution in past climates can help predict their long-term responses to climate change and give knowledge to define suitable strategies for conservation efforts (*3*, *4*).

Local adaptation, driven by natural selection, is commonly observed in plants (e.g. 45% of investigated species in (*5*). It is also prevalent in the majority of tree species, despite their long generation times (*6*). This mechanism is usually evidenced using reciprocal transplant or common garden experiments allowing to show the higher fitness of a native population in its environment of origin compared to foreign populations evolving under different environmental conditions. However, those direct approaches to study local adaptation, are feasible for short-generation plant species, while time-consuming for long-generation trees. Genome scan approaches have proven to be effective indirect methods for investigating local adaptation, either as a complement to, or independently of, common garden and reciprocal transplant experiments (*3*, *7*, *8*). Recent advances in analytical methods and Next-Generation Sequencing (NGS) technologies have enabled the use of these genome scan methods by leveraging genome-wide variation across large sample panels (*3*, *9*). Two key methodologies stand out: selective sweep detection and genomic-environment association (GEA) studies. The first method looks for local reduction of genetic diversity and/or increased linkage disequilibrium signatures linked with the rapid fixation of a beneficial mutation throughout the evolutionary history of the studied populations. These sweeps can result from adaptation to abiotic (e.g., temperature, precipitation) or biotic (e.g., pathogens) factors (*10–12*), as well as artificial selection that can eventually lead to domestication (*13*, *14*). In wild perennial species, sweeps offer valuable insights into long-term evolutionary responses to local environments (*15*). GEA approaches, on the other hand, detect statistical associations between genomic sequences and environmental variables across natural populations (*16*, *17*). This method has gained widespread application in recent years (*3*, *11*, *12*, *18*). However, successful implementation of GEA requires extensive sampling across environmental gradients and access to environmental and genomic datasets to achieve sufficient statistical power and representativeness of habitat range (*3*, *7*).

The wild olive tree, *Olea europaea* subsp. *europaea* var. *sylvestris*, is a perennial species emblematic of the Mediterranean climate (*19–21*). Its long generation time and widespread distribution along a significant latitudinal gradient make it an interesting model for studying local adaptation. The olive tree has an anemophilous and zoochorous reproduction system, combined with allogamy (*22*), leading to substantial gene flow across the MB (*23*, *24*). The evolutionary history of the olive tree in the MB dates back in the Middle Pleistocene (*25*). Since then, it has been affected by multiple glaciation events, leading to population fragmentation and persistence in climatic refugia (*26*, *27*), before significantly expanding across the MB when climatic conditions became more favourable (*28*, *29*). Deciphering this complex history, previous archaeobotanical and phylogeographical studies of the species evidenced two distinct gene-pools deriving from different glacial refuges, one in the western MB, the other in the eastern MB (*26*, *29–33*). The wild olive tree forms a genetically homogeneous wild group across several environmental regions (large climatic gradient) of the western MB, including Southern France, Corsica, Spain, and Morocco (*34*).

Around the MB, wild olive trees coexist with cultivated olive trees (*Olea europaea* subsp. *europaea* var. *europaea*) that likely mainly derived from the eastern wild gene-pool (*27*, *30*, *35–37*). Numerous hybridization events between these two genetic groups, wild and cultivated, have given rise to a genetically heterogeneous new group, the admixed group, which is widely present in natural environments alongside wild olive tree populations (*34*). The high gene flow from the cultivated compartment to the wild compartment, leading to the widespread occurrence of admixed genotypes observed in the natural environment of the western MB, raises questions about the adaptive potential of spontaneous olive populations and the selective pressures that have allowed the coexistence of both genetic groups in sympatry.

Here, we address the following three questions: (1) Are wild olive trees from the western MB locally adapted? (2) What are the major selective pressures acting on olive trees? (3) If wild olives are locally adapted, does the associated genetic variation for adaptation exist within cultivated and admixed populations? To investigate these questions, we used target sequencing data from western MB spontaneously occurring populations, cultivated olive tree accessions from western MB, as reported in Zunino et al. (2024). We examined whether local adaptation could be detected in wild olive trees using selective sweep detection and genomic-environment association (GEA) analysis. Based on the association results, we analysed SNPs allele frequencies to determine the main selective pressure, and gene annotations. Additionally, we checked whether candidate adaptive variants detected in the wild compartment could be found in the cultivated and admixed olive trees.

## Methods

### 2.1 Sampling and sequencing

A total of 504 *Olea europaea* subsp. *europaea* individuals were analysed, including 359 spontaneously occurring olive trees (*O. europaea* subsp. *europaea* var. *sylvestris*) sampled in the western MB. The sampling strategy and sequencing protocols for all accessions are detailed in Zunino et al. (2024). Briefly, the spontaneously occurring populations were collected from 26 sites across France, Spain and Morocco within the western MB (Fig. S1, Table 1). The sampling sites were selected based on previous studies of Besnard et al. (2013) (*30*) and via the Conservatoire Botanique National Simethis database (http://simethis.eu). The selection of site was also guided by their climatic data using CHELSA V2.1 database to avoid redundancy in climatic conditions at a local scale (*38*, *39*). The selection method is detailed in supporting information (Method S1, Fig. S2). These sites contained 12 to 15 spontaneous trees that can be considered as either wild or admixed olive trees (*34*). The dataset also encompassed 145 cultivated olive tree accessions from western MB (Table S1). DNA extraction, library preparation and target sequencing methods are described in Zunino *et al.* (2024). Briefly, the olive trees were sequenced using a NovaSeq^TM^ sequencing platform, with a target sequencing experiment targeting the 55,595 annotated genes of the reference genome ‘Farga’ v2 (*40*) (Table S2). We retained 210,367 baits, targeting the chromosomes and the scaffolds with a captured length of 16.8Mb. The raw sequence reads have been deposited in the European Nucleotide Archive (PRJEB61410).

**Table 1.** Summary information of the 26 spontaneously occurring populations populations of *O. europaea* subsp. *europaea* var. *sylvestris* used in this study and their uncorrelated environmental data. Climate data extracted from CHELSA V2.1 (*38*, *39*).

| Sites | Longitude | Latitude | Country | Temperature relative variables |  |  |  |  |  | Precipitation relative variables |  |  |  |
| --- | --- | --- | --- | --- | --- | --- | --- | --- | --- | --- | --- | --- | --- |
|  |  |  |  | Bio1 | Bio2 | Bio4 | Bio6 | Bio8 | Bio9 | Bio12 | Bio14 | Bio15 | Bio19 |
| F01 | 3.013 | 42.937 | France | 15.8 | 7.7 | 585 | 4.65 | 12.5 | 23.6 | 609.2 | 14.9 | 40.2 | 188.6 |
| F02 | 3.303 | 43.541 | France | 14.3 | 8.3 | 584 | 3.05 | 10.9 | 22.2 | 866.5 | 22.6 | 40.1 | 247.5 |
| F03 | 3.792 | 43.771 | France | 13.3 | 9.8 | 607 | 1.25 | 13.6 | 21.5 | 1069 | 28.6 | 42.3 | 290.6 |
| F04 | 4.733 | 43.877 | France | 15 | 9.9 | 659 | 1.75 | 15.2 | 23.9 | 705.1 | 27.1 | 40.7 | 153.5 |
| F06 | 6.361 | 43.383 | France | 15.6 | 9.2 | 624 | 2.65 | 12 | 24 | 846.4 | 17.9 | 43.1 | 232 |
| F07 | 7.303 | 43.689 | France | 15.8 | 5.4 | 538 | 5.55 | 13.2 | 22.8 | 798.1 | 16.2 | 50 | 224.1 |
| F08 | 9.064 | 42.654 | France | 16.7 | 5.3 | 545 | 6.85 | 14.3 | 23.9 | 574.8 | 9.6 | 42.7 | 153.6 |
| F09 | 8.699 | 42.404 | France | 16.6 | 4.4 | 522 | 7.45 | 14.6 | 23.4 | 1215 | 13.3 | 51.6 | 376.8 |
| F10 | 8.869 | 41.75 | France | 16.2 | 6.2 | 525 | 6.45 | 14 | 23 | 1158 | 15.9 | 48.4 | 354.2 |
| F11 | 9.202 | 41.372 | France | 16.8 | 2.7 | 471 | 9.35 | 15.4 | 22.9 | 600.7 | 7.3 | 50.5 | 162.3 |
| S12 | 3.219 | 42.235 | Spain | 15.6 | 3.7 | 479 | 7.75 | 17 | 21.9 | 644.3 | 18.2 | 38.3 | 173.9 |
| S13 | 1.976 | 41.421 | Spain | 14.9 | 7.4 | 545 | 3.75 | 16.2 | 22 | 640.1 | 25.2 | 32.2 | 129.4 |
| S14 | 0.935 | 41.02 | Spain | 16.6 | 6.3 | 535 | 6.45 | 18 | 23.6 | 561.1 | 14.8 | 43.2 | 116.7 |
| S15 | 0.386 | 40.341 | Spain | 17.8 | 8.1 | 551 | 6.15 | 18.8 | 25 | 604 | 12.5 | 41 | 147.7 |
| S16 | 0.195 | 38.803 | Spain | 17.9 | 2.9 | 463 | 10.7 | 19.8 | 23.8 | 776.6 | 6.7 | 50.7 | 218.2 |
| S17 | -3.507 | 38.374 | Spain | 16.8 | 10.8 | 719 | 2.35 | 12.3 | 26.5 | 534.3 | 4.9 | 51.2 | 183.3 |
| S18 | -6.209 | 37.256 | Spain | 18.7 | 11.1 | 571 | 5.95 | 15.4 | 26.2 | 612 | 2 | 68.8 | 248.6 |
| S19 | -3.85 | 36.766 | Spain | 17.8 | 3.8 | 428 | 10.3 | 13.7 | 23.4 | 464 | 1.2 | 68.1 | 189.5 |
| S20 | -5.669 | 36.062 | Spain | 18 | 5.3 | 368 | 10.5 | 14.5 | 22.6 | 710.3 | 1.9 | 76.5 | 326.8 |
| M21 | -5.925 | 35.79 | Morocco | 17.5 | 5 | 366 | 10.4 | 14.1 | 22.1 | 733.9 | 2.4 | 75.6 | 323.6 |
| M22 | -5.515 | 35.783 | Morocco | 17.6 | 5.9 | 381 | 9.75 | 14.1 | 22.5 | 639.7 | 1.6 | 74.2 | 288.9 |
| M24 | -5.908 | 33.534 | Morocco | 16.9 | 12.8 | 514 | 3.55 | 11.3 | 23.6 | 512.5 | 4.2 | 58.2 | 205.1 |
| M25 | -5.588 | 33.1 | Morocco | 16.4 | 12.7 | 632 | 1.65 | 8.75 | 24.8 | 719.1 | 7.6 | 62.7 | 319.3 |
| M28 | -8.04 | 31.21 | Morocco | 17.2 | 12.4 | 545 | 3.65 | 13.4 | 24.5 | 431.1 | 8.5 | 43.7 | 130.4 |
| M29 | -9.37 | 30.631 | Morocco | 18.3 | 12.4 | 539 | 5.35 | 11.6 | 25.4 | 360 | 1.7 | 82.8 | 182.4 |
| M30 | -9.691 | 31.111 | Morocco | 19.1 | 8.1 | 350 | 10.3 | 15.6 | 23.6 | 379.2 | 1.3 | 85.3 | 195.2 |

*N*= number of individuals, longitude and latitude in wgs84, Bio1: Mean annual air temperature (°C), Bio2: Mean diurnal temperature arrange (°C), Bio4: Temperature seasonality (°C/100), Bio6: Mean daily minimum air temperature of the coldest month (°C), Bio8: Mean daily air temperatures of the wettest quarter (°C), Bio9: Mean daily air temperatures of the driest quarter (°C), Bio12: Annual precipitation amount (Kg.m-2.year-1), Bio14: Precipitation amount of the driest month (kg m-2 month-1), Bio15: Precipitation seasonality (kg.m-2), Bio19: Mean monthly precipitation amount of the coldest quarter (kg.m-2.month-1).

### 2.2 SNP calling and SNP filtering

The SNP calling as well as the SNP filtering were performed as described in Zunino *et al.* (2024). After sequencing, we obtained 89,322,843 genetic and structural variants. We filtered the raw data with VCFtools version 0.1.16 (*42*). We removed indels, kept only SNPs with a quality superior to 200 and removed clusters of 3 SNPs in 10 bases. We kept only biallelic SNPs, with a minimum mean depth of 8 and maximum depth of 400. We removed sites with more than 15% missing data, then we removed individuals with more than 20% of missing data. Only the sampling sites with at least 12 individuals were considered in this study. Filtering was carried out to exclude positions with high levels of heterozygosity (>85% across all the individuals of the dataset). At this step, we obtained 316,639 SNPs spanning the whole genome.

### 2.3 Assignation of individual olive tree

We initially distinguished cultivated olive tree accessions from spontaneous occurring populations of western MB based on passport data, resulting in 145 cultivated olive trees and 359 western spontaneous occurring individuals. For our study, a second classification was performed on the western spontaneously occurring trees to differentiate wild olives from admixed olives. To separate spontaneously occurring individual olives from western MB based on their genetic background, we performed a genetic structure analysis using the sparse non-negative matrix factorization (sNMF) algorithm implemented in the LEA R package (version 3.11.3; (*43*), following the same parameters and criteria as described in Zunino et al. (2024). This analysis included all individuals: cultivated and naturally occurring populations from the western MB. For western spontaneously occurring populations, we extracted the individual ancestry coefficients assigned to a genetic ancestry cluster likely corresponding to a western wild origin as estimated by sNMF. Individuals with an assignment probability above or equal to 70% to this cluster were classified as genetically “wild”, whereas individuals below this threshold were categorized as “admixed”. We separated the genotypes according to their belonging to groups and applied within each group a filter on minor allele frequency (MAF) set to 5%. To mitigate the effects of linkage disequilibrium (LD), which could lead to the detection of false positives (*44*, *45*), we also generated a second dataset for the western wild olives with pruned SNP. To this aim, we filtered the variants with PLINK version 2.0 (*46*) to remove linked variants with a *r*^2^>0.2 in a sliding window of 50 base pairs, resulting in 71,632 unlinked SNPs for genuine wild olive trees.

### 2.3 Detection of candidate selected SNPs

The detection of selective sweeps was performed with pcadapt 4.3.3 (*47*) focusing on the individuals from the wild genetic group. The number of principal components K was chosen according to the percentage of variance explained by each PC. Significance threshold was determined either with a false discovery rate (FDR) approach with a rate set of 10% or with a *p-value* threshold at 5% corrected for multiple testing using a Bonferroni correction. As our FDR set up appeared less conservative and encompassed all the SNPs detected with the Bonferroni method, the SNPs retained according to the FDR were considered as minor outliers, among which the SNPs whose *p-values* satisfied the Bonferroni correction threshold were considered as major outliers.

### 2.4 Selection of climatic data

The environmental variables for each sampling site were extracted from the 19 Bioclimatic variables of CHELSA V2.1 from 1981 to 2010. This dataset has a spatial resolution of 30 arc seconds, corresponding to one km square at global elevation (*38*, *39*) (Table S3).

Using a set of scripts developed by Dauphin (2024) (*42*), we examined the correlations between the environmental variables in three steps: first, a principal component analysis (PCA) was performed on R version 4.2.0 Subsequently, a correlation matrix with a Pearson test was generated. When the correlation between two environmental variables was found to be higher than 75%, only one (the first to appear in the table) of the two was retained for the study. To ensure the remaining environmental variables had minimal correlation, a second PCA was performed.

### 2.5 Genomic environment association analyses

Genomic environment analysis (GEA) was conducted using a latent factor mixed-model analysis (function LFMM2 from LEA v3 package) (*49*) within the wild genetic group. An initial genetic structure analysis was carried out using sNMF from the LEA package version 3.11.3 (*50*) to determine the number of the optimal number of ancestral genetic clusters based on cross-entropy criterion. This allowed imputation of missing data in each dataset according to the best practices of LEA package for lfmm2 analysis (http://membres-timc.imag.fr/Olivier.Francois/LEA/index.htm). The lfmm2 analysis uses a two-step approach to estimate association between genetic and environmental variables. In a first step, the estimation function (lfmm2), build an object containing the latent factors (confounding factors, e.g. genetic structure) which will be included as correction factors, then in a second step, the test function (lfmm2.test) computes the significance of associations between the allelic frequencies at each SNP and the environmental variables chosen, using linear model tests. The environmental variables were tested one by one or being computed together for the test. We used the wild pruned dataset (excluding variants in linkage disequilibrium) for the estimation of latent factors, while the test function was performed using all wild variants. This allowed to reduce the possible false positive associations due to high LD in some regions. The tests were conducted 1) on all the uncorrelated environmental variables tested together (call "*All*", containing Bio1, Bio2, Bio4, Bio6, Bio8, Bio9, Bio12, Bio14, Bio15, Bio19), 2) on group of temperature-related variables (call "*Temperature*", containing Bio1, Bio2, Bio4, Bio6, Bio8, Bio9), 3) on group of precipitation-related variables (call "*Precipitation*" containing Bio12, Bio14, Bio15, Bio19) then 4) on each variable separately (Table 1). Significant outliers were assessed using either Bonferroni correction (alpha = 0.05, n= 96,648) or false discovery rate (FDR = 10%) with R package *q-value* version 2.30.0 (*49*).

### 2.6 Biological significance of candidate SNPs to adaptation

We conducted a gene functional annotation analysis within genomic regions containing outlier loci detected from both pcadapt and GEA analysis. Each gene containing associated SNPs were identified using bed windows from bedtools version 2.30.0 (*40*) to intersect the outliers with the annotated reference genome Farga V2 (*51*) considering a genomic window of 500bp upstream and downstream of the significant locus. In cases where a SNP was hitting two different genes due to geneID overlapping in the annotation of the reference genome Farga V2 (*52*), we chose to retain both. For each identified gene, we determined their potential function by assigning their specific gene ontology (GO) term and function with GO.db version 3.16.0 (*53*) and AnnotationDbi version 1.60.2 (*54*). Subsequently, we performed GO term enrichment tests for each group of outliers examined with clusterProfiler version 4.6.2 (*40*, *55*). Only the enriched GO terms with an adjusted enrichment *p-value* below 0.05 combined with a *q-value* under 0.1 were considered as significant. The results were graphically represented with the enrichplot package version 1.18.4 (*56*).

SnpEff version 4.3 was used to annotate and predict the effects of the outlier SNPs on genes and proteins. The database for SNPeff was created using the reference genome annotation, the DNA sequences, the protein sequences and the transcripts (*44*). We used the information to estimate the putative deleteriousness of each outlier which can be either high (e.g. stop gained), moderate (e.g. missense-variant), modifier (e.g. downstream gene variant) or low (e.g. synonym variant).

### 2.7 Characterization of the candidate SNPs for adaptation to environment variables and within genotypes groups

To investigate the relationship between genetic variation (represented by SNP allele frequencies) and environmental factors (represented by climatic data) among populations per site, we performed a Procrustes analysis between independent PCAs conducted on allele frequencies of candidate SNPs, identified from GEA, and the one derived from climatic variables.

Procrustes analysis is a statistical technique employed to assess the degree of similarity between two multivariate datasets by aligning their respective principal component configurations. This method involves the generation of PCAs for each dataset, then the rotation, translation, and scaling of one dataset to optimally superimpose it onto the other, thereby enabling a comparative analysis of their morphological structures. The objective is to minimize the sum of squared distances between corresponding points of the two PCA configurations. A statistically significant Procrustes fit indicates a high level of morphological congruence between the datasets, suggesting a strong correlation in their shape structures (*57*). Based on this analysis, principal component configurations have been compared for each population in each sampling site. Specifically we compared the configuration of population in the PCA based on allele frequencies of the GEA associated SNPs with the configuration of population in the PCA based on the specific sites environmental variables. A strong concordance between allele frequency patterns and climatic data would suggest the presence of strong selective constraints by the climate on this site. The frequency of alleles were determined using PLINK 2.0 --freq (*57*), Environmental variables were standardized prior to PCA analysis using the decostand function from the vegan R package (*11*, *12*, *58*, *59*). Both PCA were then aligned, the fit between the two configurations was quantified using Procrustes statistic (sum of squared residuals), then the significance of the correlation between the two PCAs was assessed using a permutation test using vegan package 2.6.4 (*60*, *61*). The test was done using outliers resulting from different GEA tests, based on different climatic datasets: *“Temperature”* related variables *“Precipitation”* related variables and “*All” variables*. When a SNP was considered as missing data for a site, it was removed from the dataset.

We further examined whether the candidate SNPs identified as outliers by GEA and/or pcadapt analyses are also present in the genomes of cultivated and admixed olive trees. Specifically, we retrieved the list of outlier SNPs, including both their genomic positions and variant identifiers. We then intersected this list with the variant datasets of cultivated and admixed individuals to determine the presence or absence of each candidate SNP in these groups. This allowed us to evaluate the extent to which cultivated and admixed olives may harbour alleles potentially involved in local adaptation to climatic conditions, thereby serving as reservoirs of adaptive genetic variation.

## 3 Results

### 3.1 Identification of wild olive genotypes in western Mediterranean Basin

The population structure analysis used to assign the 359 naturally occurring olive trees is presented in the Supporting Information (Fig. S3, Table S4). Based on this assignment, we identified 135 individuals as western wild olive genotypes and 224 as western admixed genotypes (Fig. S4). The groups of western MB wild genotypes resulting from this classification are illustrated in Figure 1.

**Figure 1.**
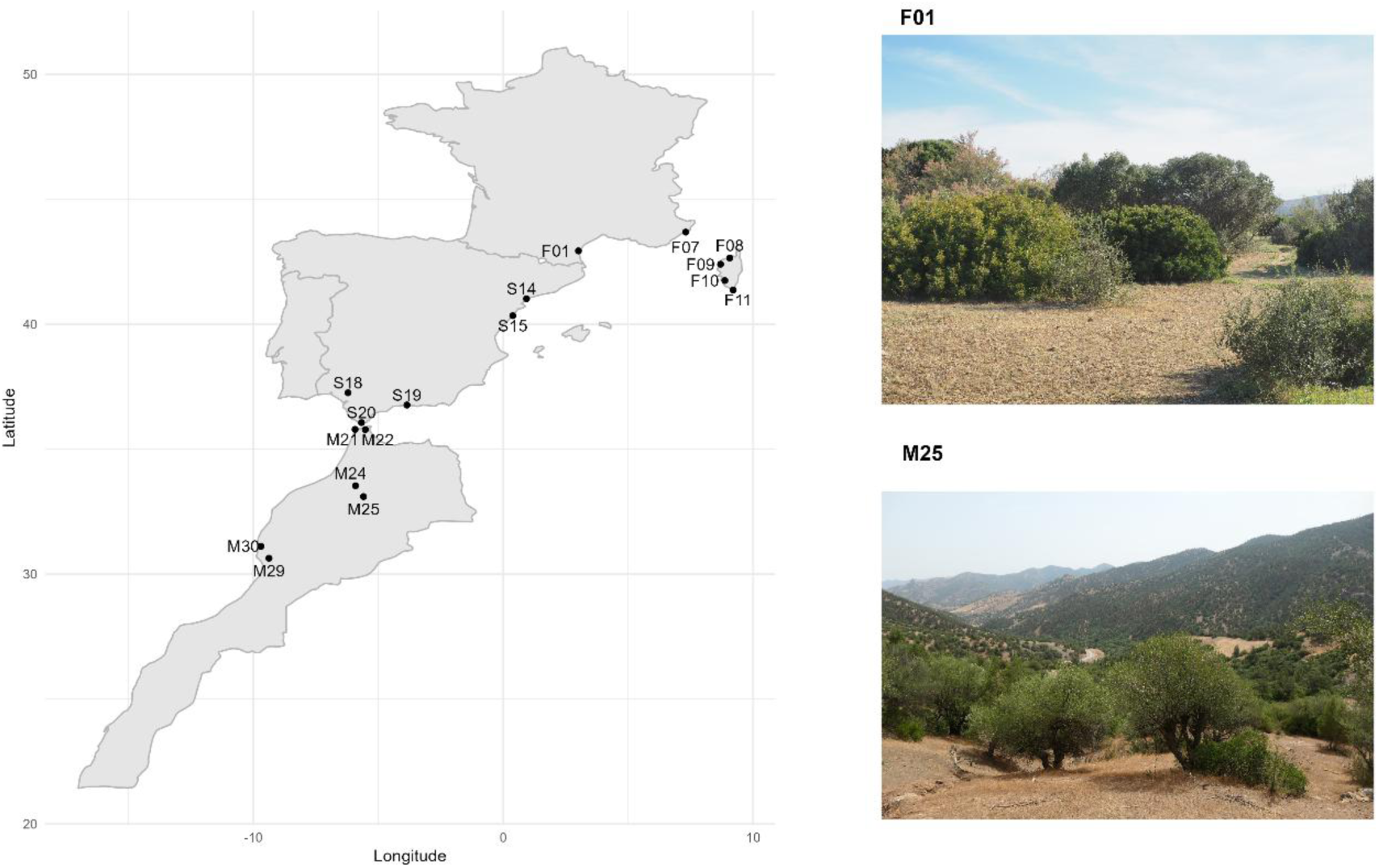
Geographical distribution of the wild population of olive trees *O. europaea* subsp*. europaea* var*. sylvestris* (n = 135) sampled in France, Spain and Morrocco. The wild olive trees were identified based on population structure with sNMF. Picture of several wild trees in F01 population, south France © Lison Zunino and in M25 population, Morrocco © Bouchaib Khadari.

### 3.2 Identification of adaptive SNPs in wild olives

The footprints of selection analysis using pcadapt detected 1,168 SNPs (Table S5). These results take into account the detection of minor outlier SNPs using a *q-value* threshold of < 0.1, including significant major outlier SNPs with the Bonferroni correction (α = 0.05, n = 96,648, adjusted threshold = 5.17e-07). Within the 1,168 SNPs, 124 were identified as major outliers (Fig. 2A, Table 2, Fig. S5 to S7).

**Figure 2.**
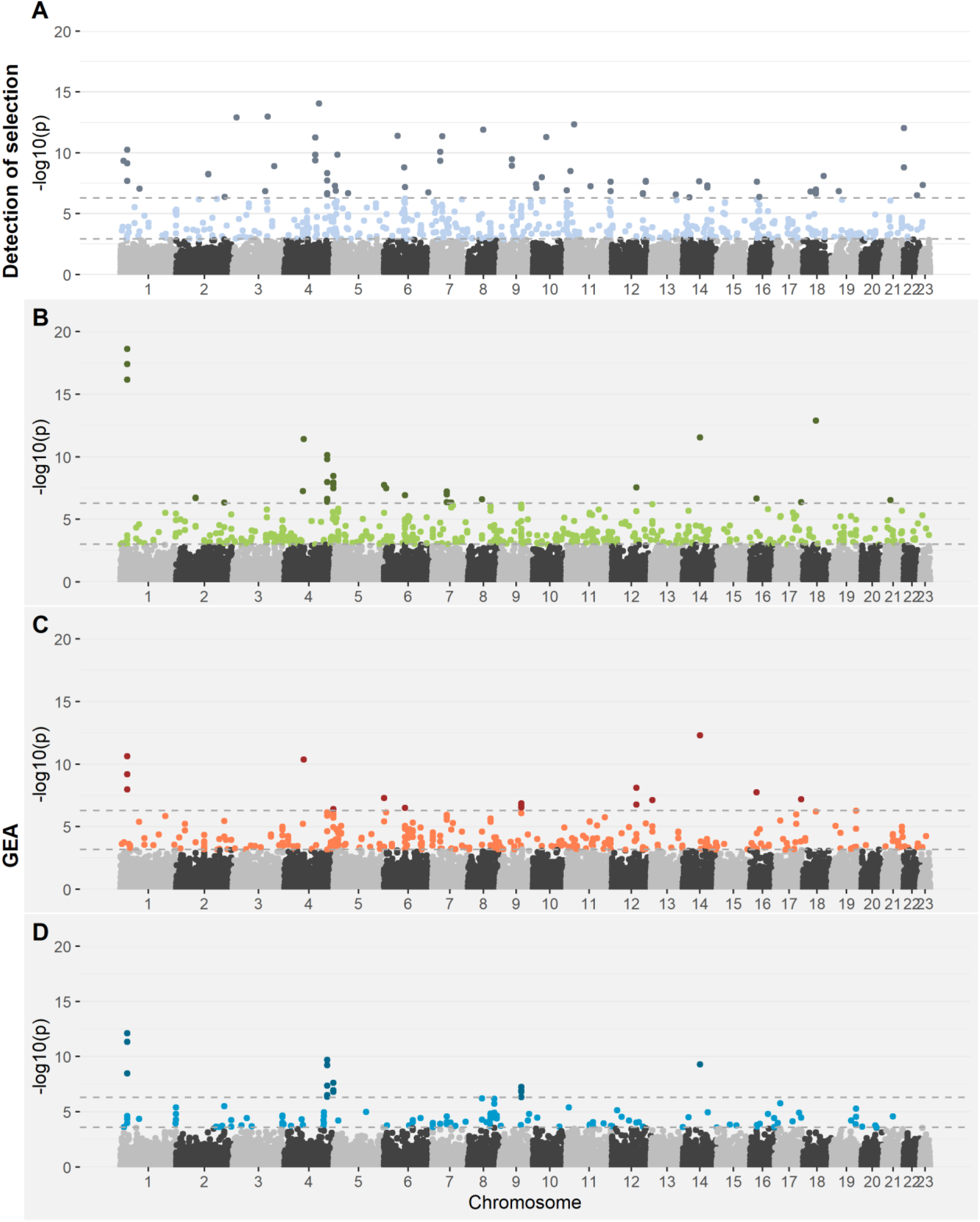
Genome-wide scans for selective sweep detection and genome-environment associations (GEAs) on chromosome. Colours indicate the importance of the outlier SNP, with dark shades representing ‘major’ outliers (*p-value* < 0.05 with Bonferroni correction–alpha =0.05, n=96,648, adjusted threshold = 5.17e-07) and light shades representing ’minor’ (*q-value* < 0.1, excluding ‘major’). (A) results of selective sweep using pcadapt, (B) results of environmental association to “All” related environmental variable with LFMM (C) results of environmental association to ‘Temperature” related variable using LFMM (D) results of environmental association to “Precipitation” related variable using LFMM.

**Table 2.**
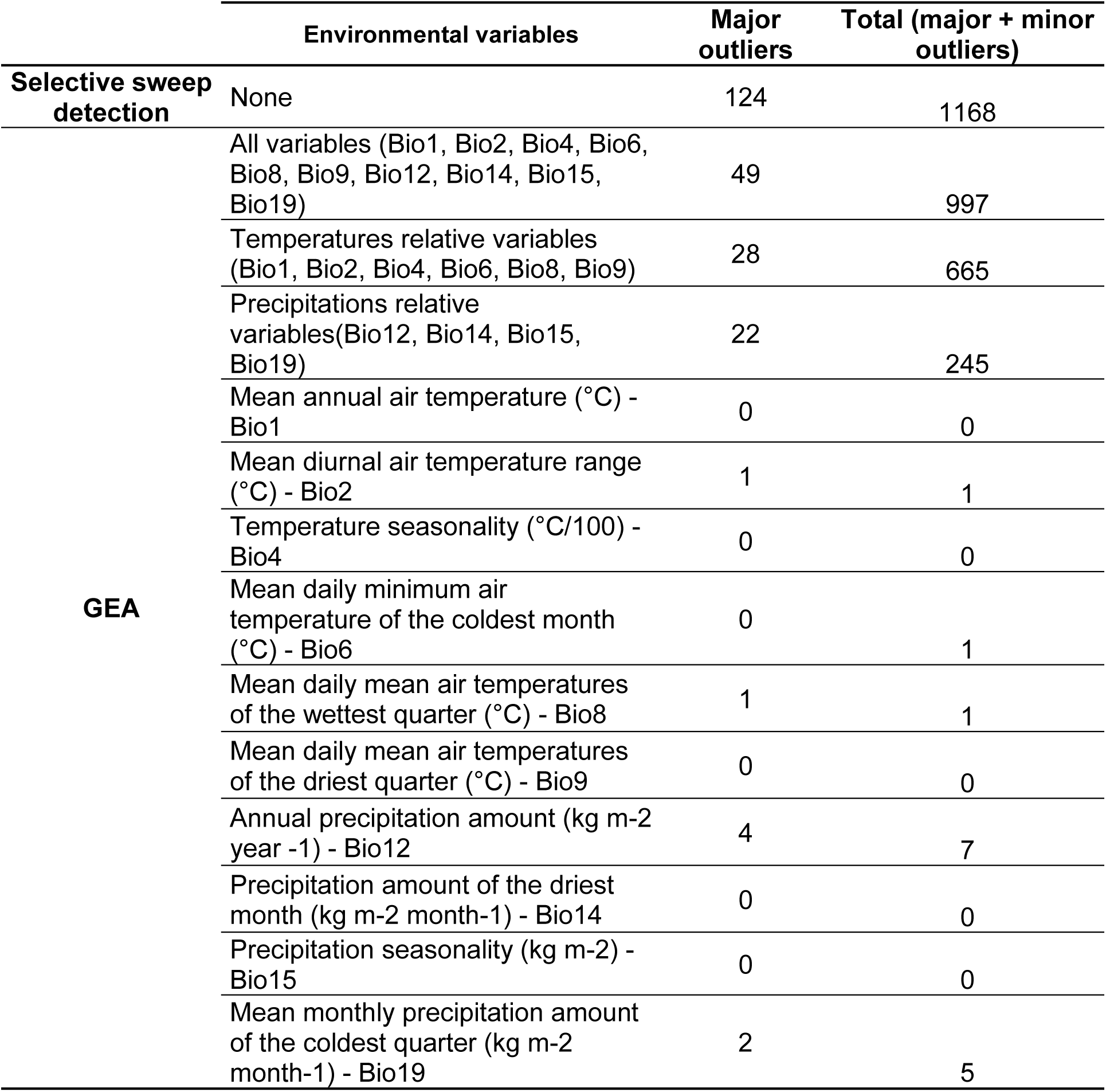
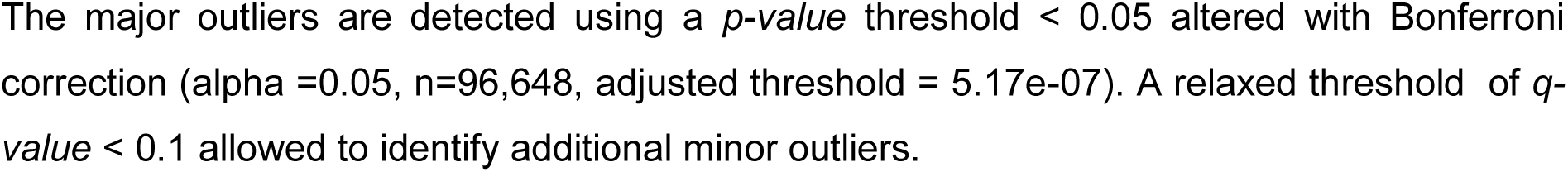
Number of candidate SNPs for selection resulting from pcadapt analysis (selective sweep detection) and for association to climatic variables resulting from LFMM analysis for each environmental variable (Genomic environment association analysis).

The GEA was performed using the following selected uncorrelated environmental variables : *mean annual air temperature* (Bio1), *mean diurnal air temperature range* (Bio2), *temperature seasonality* (Bio4), *mean daily minimum air temperature of the coldest month* (Bio6), *mean daily mean air temperatures of the wettest quarter* (Bio8), *mean daily mean air temperature of the driest quarter* (Bio9), *annual precipitation amount* (Bio12), *precipitation amount of the driest month* (Bio14), *precipitation seasonality* (Bio15) and *mean monthly precipitation amount of the coldest quarter* (Bio19) (Table 1, Fig. S8, Fig. S9, Fig. S10). The results for wild olive were considered as significant and major outliers when *p-values* < 5.17e-07. The FDR threshold for considering association as significant varied depending on tested environmental variable between 5.12e-08 and 1.03e-03 (Table S6).

The wild olive trees genetic group revealed 997 SNPs in total selected by “*All*” including 49 major and 948 minors outliers. There were 665 outliers in total related to “*Temperature*”, including 28 major and 637 minor outliers. Regarding the group of variables related to “*Precipitation*”, 245 associated SNPs were found, including 22 major outliers (Fig. 2B to 2C, Table 2, Table S7). By looking for individual environmental variables separately, SNPs were selected by five environmental variables. Three temperature variables have associated outliers: one major outlier for "*Bio2*", one minor outlier for "*Bio6*" and one major outlier for “*Bio8*”. For precipitation, there were also two variables with associated SNPs: seven outliers in total including four majors for "*Bio12*”, and five outliers in total for "*Bio19*", including two majors (Table 2, Fig. S11 to Fig. S13).

### 3.3 Association between climatic gradients and outlier allele frequencies

Procrustes tests comparing PCAs of outlier SNP frequencies, obtained from different GEA analyses and the PCA based on environmental variables across populations were performed. SNPs associated with Bio6, Bio8, and Bio2 were excluded from the analysis due to the presence of missing data for at least one sampling site. As a result, these climatic variables were not tested using Procrustes analysis. The complete list of SNPs retained for each analysis is provided in Supporting information (Table S8). Each remaining SNPs detected according to GEA, were tested against environmental data (Target PCA): “*All*” climatic data, “*Temperature*” related data, “*Precipitation*” related data (Fig. 3). When looking at Target PCA “*All*” climatic data, significant associations were found for the outliers SNPs of "*All*", "*Temperature*" related variables, "*Precipitation*" related variables, Bio12 and Bio19, with global Procrustes distances (m12^2^) ranging from 0.679 to 0.8056. Low m12^2^ values indicate stronger correspondence between the genetic and environmental datasets. Among the tested PCA of “All” climatic data, the tested PCA of outliers SNPs frequencies detected by “*Precipitation*” related variables showed the strongest match and correlation (m12^2^ = 0.679, cor = 0.5666, p = 0.002, permutations = 999), followed by outliers SNPs frequencies detected by Bio19 (m12^2^ = 0.7144, cor = 0.5344, p = 0.003, permutations = 999), the outliers SNPs frequencies detected by "*All*" climatic variables (m12^2^ = 0.7388, cor = 0.5111, p = 0.005, permutations = 999), and the outliers SNPs frequencies detected by "*Temperature*" (m12^2^ = 0.7623, cor = 0.4876, p = 0.023, permutations = 999) and the outliers SNPs frequencies detected by Bio12 (m12^2^ = 0.8056, cor = 0.4409, p = 0.045, permutations = 999). For the target PCA of “*Temperature*” related data, only one tested PCA, outliers SNPs frequencies detected by “*Precipitation*” related variables is significant (m12^2^ = 0.7832, cor = 0.4656, p = 0.038, permutations = 999). In contrast, Target PCA of “*Precipitation”* related data exhibited significant correlations with all Tested PCA. The strongest Procrustes fit was observed for outliers SNPs frequencies detected by Bio19 (m12^2^ = 0.4742, cor = 0.7251, p = 0.001, permutations = 999), followed by the outliers SNPs frequencies detected by "*Precipitation*" related variables (m12^2^ =0.5249, cor = 0.6892, p = 0.001, permutations = 999), the outliers SNPs frequencies detected by Bio12 (m12^2^ = 0.6224, cor = 0.6145, p = 0.001, permutations = 999), the outliers SNPs frequencies detected by "*All*" climatic variables (m12^2^ = 0.65, r = 0.5916, p = 0.005, permutations = 999), and finally the outliers SNPs frequencies detected by "*Temperature*" related variables (m12^2^ = 0.739, cor = 0.5109, p = 0.02, permutations = 999) (Fig.3).

**Figure 3.**
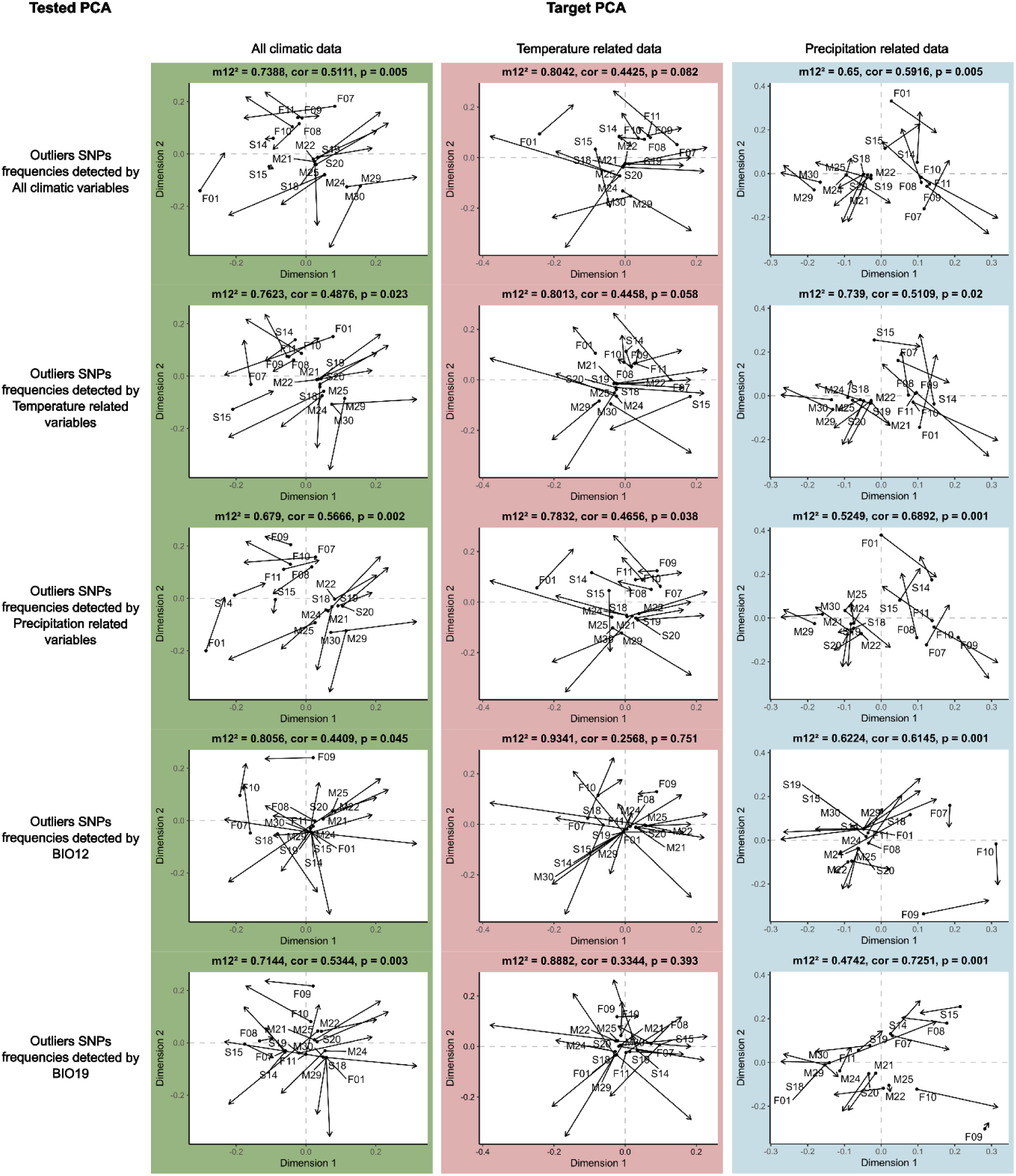
Comparison between PCA obtained from allelic variation of GEA outlier SNPs (Tested PCA) and PCA obtained from climatic variation (Target PCA). The Tested PCA from top to bottom is considering the outliers SNPs frequencies detected by GEA for all climatic variables, temperature related variables, precipitation related variables, Bio12 and Bio19. The Target PCA is considering “All” climatic data (green plots), the “Temperature” related data (red plots) or the “Precipitation” related data (blue plots) using Procrustes test. The black arrows indicate the transformations needed to align the spatial configuration of outliers SNP frequency per population to the climatic data per population, after rotation to maximum similarity between the two ordinations. The m12² (Procrustes sum of squared residuals) is the percent correspondence between the two ordinations, cor is the correlation coefficient and p the *p-value* from permutation tests.

### 3.4 Comparison of outlier detection between selective sweep detection and GEA analyses

We also compared the results of pcadapt and GEA analysis. We detected 204 unique common variants, associated with 143 unique genes (Table 3). Of these 204 variants, 130 were identified in the results of the "*Temperature*" related climate GEA test, 85 in the "*Precipitation*" related climate test, three in the *Bio12* GEA test, and two in the *Bio19* GEA test.

**Table 3.** Unique gene annotations and outliers SNPs across two detection methods, pcadapt and LFMM. Common refers to genes or variants detected in both analyses.

| Identification methods | Unique variants | Annotated corresponding genes |
| --- | --- | --- |
| pcadapt | 1168 | 735 |
| LFMM | 1205 | 757 |
| Overlap-common | 204 | 143 |

### 3.5 Functional annotation and characterisation of adaptive SNPs in wild olives

Among the unique SNPs identified using both pcadapt (1,168 SNPs) or GEA (1,205 SNPs), a large proportion were associated with annotated genes based on the structural genome annotation of the reference genome *Farga V2* (Julca et al. 2020). Specifically, 1,114 SNPs from the pcadapt analysis were linked to 735 distinct genes, while 1,125 SNPs from the GEA analysis were associated with 757 genes (Table 3). In both cases, some SNPs were mapped to multiple genes due to overlapping gene models in the genome annotation, in which case the different annotation were retained.

Functional annotation resulted in 571 functionally annotated genes for the candidate SNPs from pcadapt analysis (Table S5) and 601 functionally annotated genes and their orthologous functions for the candidate SNPs GEA analyses (Table S7). Gene Ontology (GO) enrichment analysis revealed no significant enrichment for candidate SNPs identified by pcadapt. In contrast, five significantly enriched GO terms were detected among the candidate SNPs identified by GEA, including four unique go-terms: “callose localisation” (GO:0052545; *p-value* 1.86e-05 and 1.46e-04 respectively), which appeared in both in the “*All*” candidate SNPs and the “*Temperature*” related variables candidate SNPs, and three terms exclusively found in the “*All”* candidate SNPs subset, “reactive oxygen species metabolic process” (GO:0072593; 1.51e-04), “response to ozone” (GO:0010193;3.00e-04), and “phosphatidylinositol-4,5-bisphosphate binding” (GO:0005546; 4.38e-04) (Fig. S14). Predicted functional impacts of the SNPs, assessed using SNPeff, showed that the majority were classified as having a modifier effect, 89% for pcadapt and 73% for GEA, typically corresponding to exon variants or variants in downstream regions. High-impact variants were rare: one SNP from the pcadapt analysis was predicted to cause a ‘stop gained’ mutation affecting the gene OE9A021200 involved in various function as in microtubule cytoskeleton organization or response to salt stress (Table S9), while 0.6% of GEA candidates SNPs were classified as high-impact (e.g., stop gained, start lost, stop lost, splice region or intron variants), involved in different functions (Table S9).

Given the results from the Procrustes analysis, which highlighted the importance of SNPs identified through the GEA approach in association with the bioclimatic variables BIO12 and BIO19, we conducted a functional annotation of the sequences surrounding these loci. Specifically, we retrieved annotation sequences spanning 1 kilobase (kb) upstream and downstream of the seven SNPs associated with BIO12 and the five SNPs associated with BIO19. These sequences were subjected to BLAST analysis to infer potential gene function. The SNPs linked to BIO12 were predominantly associated with genes encoding Annexin D1 and Annexin GH1, as well as a conserved hypothetical protein. For BIO19-associated SNPs, functional annotations indicated putative involvement in Regulatory protein MLP, WLIM1-like protein, Pyruvate decarboxylase 2, and conserved hypothetical protein (Table S10).

### 3.6 Occurrence of environment-associated SNPs in cultivated and admixed olives

We compared the SNPs outliers obtained from both approaches, pcadapt and GEA, with cultivated and admixed SNP datasets. Out of the 1,168 candidate SNPs detected with pcadapt, 677 were found present within the cultivated and 913 within the admixed dataset, respectively. For the GEA, out of the 1,205 candidate SNPs detected, 590 SNPs could be found within the cultivated and 981 SNPs within the admixed dataset. By examining candidate SNPs identified in wild olive trees through GEA analyses and looking for their presence-absence patterns in cultivated and admixed olive trees, we observed that only a few SNPs were shared when considering unique environmental variable (Bio2, Bio6, Bio8, Bio12 and Bio19). In admixed olive trees, a single SNP was shared for each of *Bio2*, *Bio6*, *Bio8*, and *Bio12*, while no SNPs were shared for *Bio19*. In cultivated olive trees, three SNPs were found for *Bio12*, whereas no SNPs were detected for the other individual variables. When considering SNPs across all environmental variables ("*All*" category), 473 SNPs were present in cultivated olive trees, while 798 SNPs were found in admixed olive trees. For “*Temperature*” related variables, 369 SNPs were found in cultivated olive trees and 590 SNPs in admixed olive trees. Regarding “*Precipitation*” related variables, 119 SNPs were identified in cultivated olive trees, whereas 198 SNPs were observed in admixed olive trees (Table 4).

**Table 4.**
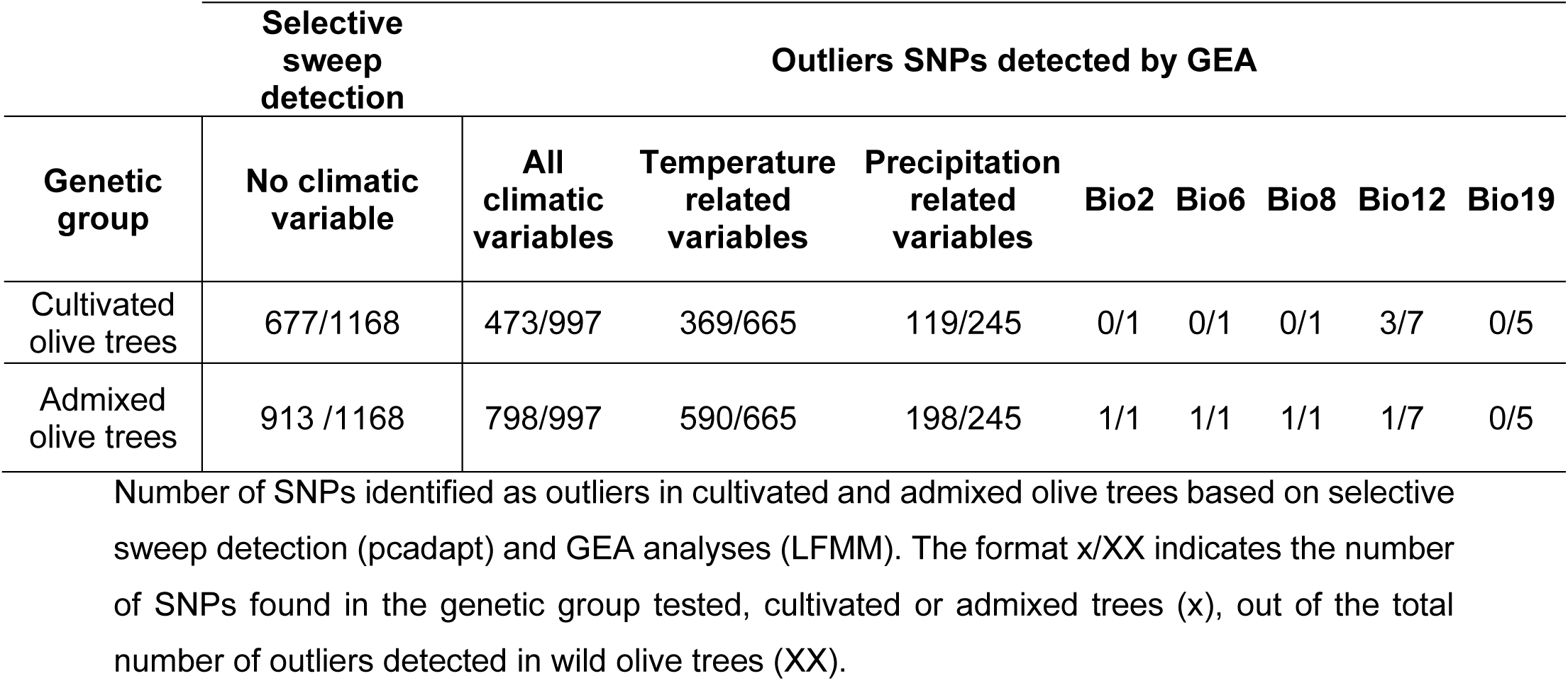
Occurrence of candidate SNPs, resulting from pcadapt and GEA analysis on wild olive trees, among cultivated and admixed olive trees dataset.

## 4 Discussion

We investigated the local adaptation in wild populations of *O. europaea* from the western Mediterranean Basin using target sequencing data across a broad latitudinal gradient. First, we conducted genome-wide scans for selection, followed by GEA analyses based on bioclimatic variables characterizing each sampling site. These analyses revealed a large number of SNPs associated with environmental gradients. We explored the likely drivers of selection for these candidate SNPs and then assessed if these SNPs were present or absent in admixed and cultivated genomes to evaluate the extent of introgression and the potential contribution of wild alleles to cultivated genotypes. We found that all climate variables have influence on population structure, with temperature-related variables being major drivers, alongside precipitation related variables such as BIO12 (annual precipitation) and BIO19 (precipitation of the coldest quarter). These findings are consistent with patterns observed in other tree species (*11*, *12*, *58*, *59*) and align well with the ecological physiology of the olive tree (*62*, *63*). A subset of the candidate variants identified in wild populations by either pcadapt and GEA analysis was also found in cultivated genotypes, and even in greater representation in admixed genotypes, raising questions about the possibility of adaptive introgression from wild olives into cultivated populations, potentially enhancing genetic diversity and adaptive potential in admixed groups.

### 4.1 Important selective pressure on western wild olives trees in natural environment drives local adaptation to climate

Detection of selective sweeps indicated that wild olive trees (*Olea europaea* subsp. *europaea* var. *sylvestris*) have undergone strong selective pressures in the western MB. Genome–environment association analyses further revealed that multiple SNPs were significantly correlated with environmental variables, by looking either at environmental variables tested all together, temperature-related variables or precipitation-related variables. When considering single variables, five of them significantly shaped the genomic diversity of the wild olive trees*: Mean diurnal air temperature range* (°C - BIO2), *Mean daily minimum air temperature of the coldest month* (°C - BIO6), *Mean daily mean air temperatures of the wettest quarter* (°C - BIO8), *Annual precipitation amount* (kg m-2 year -1 - BIO12), and *Mean monthly precipitation amount of the coldest quarter* (kg m-2 month-1 - BIO19). A small proportion of SNPs (approximately 17%) identified through GEA overlapped with those detected by selective sweep analyses. This overlap might highlight highly relevant climate-related adaptive SNPs.

Among the SNPs identified as shared between GEA and selective sweep detection results, most were associated with temperature-related variables, highlighting temperature as a potentially strong selective force shaping the genomic variation in wild olive populations across the region. This finding aligns with the broader results of our analysis as when all environmental variables were considered together, SNPs associated with temperature-related factors were the most prevalent, followed by those linked to precipitation-related variables. The concordance in the multivariable space between the structure of the climatic variables and different sets of candidate SNPs, further supports the significant role of temperature in shaping the spatial genetic structure of these populations based on allele frequencies. Impact of temperature-related variables have been previously highlighted in other species, such as eucalyptus, red spruce, mediterranean oaks and conifers (*11*, *12*, *58*, *59*). This impact of temperature appears to be strongly supportive of the temperature requirements of olive trees. Indeed, wild olive trees are found in the Mediterranean climate, characterized by specific temperature and precipitation patterns. This climate is defined by a dry summer season, which can vary from warm to hot depending on the region (*60*, *61*, *64*). Olive tree is considered as one of the main representative species of this climate (*60*). When considering olive tree biology, the impact of temperature seems very supportive, temperatures variations within the range of 0 to -3°C result in slight damage on the young tree whereas temperatures falling below -10°C can cause severe damage (*65*). The impact of low temperature tolerance has been assessed in cultivated olive tree accession showing significant variations in cold tolerance among them (*66*). Similar evaluations in wild populations have not been conducted yet, but can imply the potential for substantial variation among wild populations as well.

Concerning precipitation-related variables, the multivariate space analysis between the structure of the climatic variables and different sets of candidate SNP frequency indicates that precipitation may also play a substantial role in determining the spatial genetic structure of wild olive populations. Among the key climatic predictors associated with SNPs, the most influential were the *mean monthly precipitation of the coldest quarter* (BIO19), followed by *annual precipitation amount* (BIO12). BLAST analysis of sequences harbouring these SNPs revealed two candidate genes with known roles in plant adaptation to climatic stress. The first gene encodes a Major Latex-like Protein (MLP), a protein family known for its involvement in drought and salt stress responses, as well as pathogen resistance (*61*, *67*). The second candidate, Pyruvate Decarboxylase 2 (PDC2) has been found to be involved in plant growth development under hypoxia. The association of SNPs within PDC2 with BIO19 suggests that soil water saturation or fluctuating water regimes may drive selective pressures on olive trees. Consequently, trees carrying specific variants of these genes may exhibit enhanced tolerance to hypoxia-like stress conditions (*58*). Considering the biology of the olive tree, particularly its chilling requirements during the coldest months and its adaptation to the highly seasonal precipitation regimes characteristic of the MB, marked by wet winters and dry to nearly arid summers, these findings are ecologically meaningful (*68*). Precipitation patterns, together with temperature, emerge as important environmental drivers of selection in olive tree populations. Such selective pressures are consistent with those observed in other perennial Mediterranean species. For example, in Mediterranean conifers, precipitation and temperature have been identified as key factors shaping local adaptation (*69*). Similarly, in oleaceous species like *Forsythia suspensa* (weeping forsythia), environmental drivers such as waterlogging, cold, and drought have been shown to significantly contribute to adaptive differentiation (*70*, *71*). These parallels highlight the central role of hydric stress, either with water excess (e.g., waterlogging) or deficit (e.g., drought), along with temperature constraints, in driving local adaptation among long-lived woody plants in Mediterranean ecosystems.

Among the 1,205 candidate SNPs under selection, GO enrichment analysis identified the term ’callose localisation’ as significantly enriched (appearing twice), a process widely recognized for its role in plant responses to both abiotic and biotic stresses (*72*). The reactive oxygen species metabolic process is also well-known for its implication in plant responses to abiotic stresses, e.g. drought and salinity (*73*).

### 4.2 Differential local adaptation between wild and admixed olive trees: a potential help to cope with climate change?

By investigating patterns of selection and local adaptation information of the wild in cultivated and admixed olive genomes, we found that between 80% and 84% of genetic variants identified as putatively adaptive in wild populations are present in admixed genomes, while only 48% to 55% are retained in cultivated genomes. These findings suggest that admixed individuals maintain a higher proportion of adaptive SNPs originally identified in wild olive trees than their cultivated counterparts. Given that hybridization events likely occurred with closely related wild populations harboring environmentally adaptive alleles, admixed individuals may have inherited some degree of indirect local adaptation. However, due to the recent nature of these hybridization events, it remains uncertain whether this genetic contribution has translated into effective adaptation. Nonetheless, this introgression from wild gene pools may enhance the adaptive potential of the admixed group by increasing standing genetic variation. The retention of alleles from both wild and cultivated sources potentially allows for the presence of multiple alleles at key loci, which could facilitate a more rapid evolutionary response to environmental change (*74*, *75*). Gene flow can facilitate adaptive evolution in new environments by increasing genetic variation or fitness (*76*). These results align with the assumption that long-lived woody perennials species tend to hybridize more frequently, contributing to the high levels of genetic diversity and helping to colonize new habitats (*77*). Recent studies further support the idea that hybrids can adapt more rapidly to novel environments than their parental populations (*78*).

The significant presence of admixed individuals within naturally occurring populations, retaining a considerable portion of adaptive variants from wild lineages, support the assumption made by Lavee & Zohary (2011) (*77*) that wild olive trees, because of gene flow with the cultivated group, are led to the loss of their specific characteristics, developed over decades by natural selection, with a reduction of wild genetic diversity. Recognizing wild olive trees as a species deserving protection would allow for the implementation of conservation strategies. The implementation of these conservation measures for the wild olive populations is crucial for preserving its genetic diversity and ecological function. In-situ conservation strategies are essential to prevent the ongoing destruction of remaining natural populations from human activities. Complementary to this, ex-situ approaches, such as the establishment of a comprehensive collection of wild olive genotypes, are necessary to safeguard the species’ genetic variation across its diverse ecological range. These collections can serve as valuable reservoirs for further investigations, future restoration and breeding programs. The presence of adaptive alleles, for example, to warmer temperatures, could facilitate the establishment of these wild populations through assisted migration in future ecological niches corresponding to their current climatic conditions. Furthermore, predictive studies incorporating genomic offset analyses in conjunction with ecological niche modeling (*78*), are vital to anticipate the potential responses of wild and admixed olive trees to future climate scenarios. Such integrative approaches can help identify which genotypes are most likely to persist under changing environmental conditions, and in which types of ecological niches, thereby informing and optimizing both in-situ and ex-situ conservation planning.

## Conclusion and perspectives

In conclusion, we provided evidence that western genuine wild olive trees, *Olea europaea* var. *sylvestris* is locally adapted to its environment, mainly due to climatic temperature factors and precipitation amount per month of coldest quarter. Furthermore, admixed individuals retained at least 80% of local adaptation information in their genome, the SNPs detected as locally associated to climate in wild genetic groups. The broadening of standing genetic variation resulting from this hybridization could enable a faster response to adaptation to new or changing environments. Our study claims for a more in-depth analysis of adaptive introgression in the cultivated accessions from the western wild gene pool. This could be achieved by generating extensive Whole Genome Sequencing data for both cultivated and wild accessions and using statistical approaches such as local ancestry inferences. This study also paves the way to the prediction of future maladaptation of natural olive trees populations within the western MB, and calls for its conservation as well as assessing its importance for cultivation.

## Authors contributions

PC, BK and LZ conceptualized the project. AEB, BK and LZ collected the biological resources. PM, GS, BK, PC and LZ developed the methodology. GS and LZ did the data curation. BK led the administration project. Supervision was led by GS, EC, BK and PC. LZ led the formal analysis with input from GD, AS and PC. LR and BR participated in the interpretation of some results. LZ wrote the manuscript. The final manuscript and review were done with all the co-authors.

## Data availability

All raw sequences of Olea europaea are available in the following database: ClimOliveMed; 2023; GenomiCOM: ClimOliveMed Genomic resources for research on adaptation of olive tree to climate change; European Nucleotide Archive; 2023-04-17; PRJEB61410. Snakemake workflow of the SNP calling is available here: https://forgemia.inra.fr/gautier.sarah/ClimOlivMedCapture.

## Acknowledgements

This work has been realized with the support of MESO@LR-Platform at the University of Montpellier, the CIRAD, and the technical support of the bioinformatics group of the UMR AGAP Institute, member of the French Institute of Bioinformatics (IFB)—South Green Bioinformatics Platform.

LZ was funded by a PhD scholarship from the French government. This study was funded through Labex AGRO 2011-LABX-002, project ClimOliveMed n° 2003-001 (under I-Site Muse framework) coordinated by Agropolis Fondation. The funders had no role in study design, data collection and analysis, decision to publish, or preparation of the manuscript.

Authors are grateful to Thibaut Leroy, Olivier François, Benjamin Dauphin, Laurène Gay and Laurence Desprès for helpful comments on an early version of the manuscript.

## References

1. P. Lionello, L. Scarascia, The relation between climate change in the Mediterranean region and global warming. *Regional Environ*. Change 18, 1481– 1493 (2018).

2. F. Medail, P. Quezel, Hot-Spots Analysis for Conservation of Plant Biodiversity in the Mediterranean Basin. Ann. Mo. Bot. Gard. 84, 112–127 (1997).

3. D. A. Steane, B. M. Potts, E. McLean, S. M. Prober, W. D. Stock, R. E. Vaillancourt, M. Byrne, Genome-wide scans detect adaptation to aridity in a widespread forest tree species. Mol. Ecol. 23, 2500–2513 (2014).

4. A. Hoffmann, P. Griffin, S. Dillon, R. Catullo, R. Rane, M. Byrne, R. Jordan, J. Oakeshott, A. Weeks, L. Joseph, P. Lockhart, J. Borevitz, C. Sgrò, A framework for incorporating evolutionary genomics into biodiversity conservation and management. Climate Change Responses 2 (2015).

5. R. Leimu, M. Fischer, A meta-analysis of local adaptation in plants. PLoS One 3, 1–8 (2008).

6. F. J. Alberto, S. N. Aitken, R. Alía, S. C. González-Martínez, H. Hänninen, A. Kremer, F. Lefèvre, T. Lenormand, S. Yeaman, R. Whetten, O. Savolainen, Potential for evolutionary responses to climate change - evidence from tree populations. Glob. Chang. Biol. 19, 1645–1661 (2013).

7. S. Hoban, J. L. Kelley, K. E. Lotterhos, M. F. Antolin, G. Bradburd, D. B. Lowry, M. L. Poss, L. K. Reed, A. Storfer, M. C. Whitlock, Finding the Genomic Basis of Local Adaptation: Pitfalls, Practical Solutions, and Future Directions. Am. Nat. 188, 379–397 (2016).

8. V. L. Sork, Genomic Studies of Local Adaptation in Natural Plant Populations. J. Hered. 109, 3–15 (2018).

9. T. Capblancq, K. Luu, M. G. B. Blum, E. Bazin, Evaluation of redundancy analysis to identify signatures of local adaptation. Mol. Ecol. Resour. 18, 1223–1233 (2018).

10. T. Mitchell-Olds, J. H. Willis, D. B. Goldstein, Which evolutionary processes influence natural genetic variation for phenotypic traits? Nat. Rev. Genet. 8, 845– 856 (2007).

11. R. Jordan, A. A. Hoffmann, S. K. Dillon, S. M. Prober, Evidence of genomic adaptation to climate in Eucalyptus microcarpa: Implications for adaptive potential to projected climate change. Mol. Ecol. 26, 6002–6020 (2017).

12. T. Capblancq, S. Lachmuth, M. C. Fitzpatrick, S. R. Keller, From common gardens to candidate genes: exploring local adaptation to climate in red spruce. New Phytol. 237, 1590–1605 (2023).

13. S. I. Wright, I. V. Bi, S. G. Schroeder, M. Yamasaki, J. F. Doebley, M. D. McMullen, B. S. Gaut, The effects of artificial selection on the maize genome. Science 308, 1310–1314 (2005).

14. N. Duan, Y. Bai, H. Sun, N. Wang, Y. Ma, M. Li, X. Wang, C. Jiao, N. Legall, L. Mao, S. Wan, K. Wang, T. He, S. Feng, Z. Zhang, Z. Mao, X. Shen, X. Chen, Y. Jiang, S. Wu, C. Yin, S. Ge, L. Yang, S. Jiang, H. Xu, J. Liu, D. Wang, C. Qu, Y. Wang, W. Zuo, L. Xiang, C. Liu, D. Zhang, Y. Gao, Y. Xu, K. Xu, T. Chao, G. Fazio, H. Shu, G. Y. Zhong, L. Cheng, Z. Fei, X. Chen, Genome re-sequencing reveals the history of apple and supports a two-stage model for fruit enlargement. Nat. Commun. 8 (2017).

15. D. B. Neale, A. Kremer, Forest tree genomics: growing resources and applications. Nat. Rev. Genet. 12, 111–122 (2011).

16. V. L. Sork, S. N. Aitken, R. J. Dyer, A. J. Eckert, P. Legendre, D. B. Neale, Putting the landscape into the genomics of trees: Approaches for understanding local adaptation and population responses to changing climate. Tree Genet. Genomes 9, 901–911 (2013).

17. C. Rellstab, F. Gugerli, A. J. Eckert, A. M. Hancock, R. Holderegger, A practical guide to environmental association analysis in landscape genomics. Mol. Ecol. 24, 4348–4370 (2015).

18. T. Leroy, J. M. Louvet, C. Lalanne, G. Le Provost, K. Labadie, J. M. Aury, S. Delzon, C. Plomion, A. Kremer, Adaptive introgression as a driver of local adaptation to climate in European white oaks. New Phytol. 226, 1171–1182 (2020).

19. P. Ozenda, Sur les étages de végétation dans les montagnes du bassin méditerranéen. Documents de cartographie écologique XVI, 1–32 (1975).

20. S. Rivas Martínez, J. M. Gandullo Gutiérrez, J. L. Allué Andrade, J. L. Montero de Burgos, J. L. González Rebollar, Memoria Del Mapa de Series de Vegetación de España 1:400.000 (1987).

21. D. Kaniewski, E. Van Campo, T. Boiy, J. F. Terral, B. Khadari, G. Besnard, Primary domestication and early uses of the emblematic olive tree: Palaeobotanical, historical and molecular evidence from the Middle East. Biol. Rev. Camb. Philos. Soc. 87, 885–899 (2012).

22. P. Saumitou-Laprade, P. Vernet, X. Vekemans, S. Billiard, S. Gallina, L. Essalouh, A. Mhaïs, A. Moukhli, A. El Bakkali, G. Barcaccia, F. Alagna, R. Mariotti, N. G. M. Cultrera, S. Pandolfi, M. Rossi, B. Khadari, L. Baldoni, Elucidation of the genetic architecture of self-incompatibility in olive: Evolutionary consequences and perspectives for orchard management. Evol. Appl. 10, 867–880 (2017).

23. D. H. R. Spennemann, L. Richard Allen, The avian dispersal of olives Olea europaea: Implications for Australia. Emu 100, 264–273 (2000).

24. H. Ando, V. Martín-Vélez, G. Tavecchia, A. Traveset, I. Jiménez-Martín, J. M. Igual, A. Martínez-Abraín, S. Hervías-Parejo, Gulls contribute to olive seed dispersal within and among islands in a Mediterranean coastal area. J. Biogeogr. 51, 131–139 (2024).

25. P. U. Clark, A. S. Dyke, J. D. Shakun, A. E. Carlson, J. Clark, B. Wohlfarth, J. X. Mitrovica, S. W. Hostetler, A. M. McCabe, The Last Glacial Maximum. Science 325, 710–714 (2009).

26. Y. Carrión, M. Ntinou, E. Badal, Olea europaea L. in the North Mediterranean Basin during the Pleniglacial and the Early-Middle Holocene. Quat. Sci. Rev. 29, 952–968 (2010).

27. G. Besnard, B. Khadari, M. Navascués, M. Fernández-Mazuecos, A. E. Bakkali, N. Arrigo, D. Baali-Cherif, V. Brunini-Bronzini de Caraffa, S. Santoni, P. Vargas, V. Savolainen, The complex history of the olive tree: From late quaternary diversification of mediterranean lineages to primary domestication in the northern Levant. Proceedings of the Royal Society B: Biological Sciences 280, 1–7 (2013).

28. N. Planchais, Palynologie lagunaire: l’exemple du Languedoc-Roussillon. Annales de Géographie 93, 268–275 (1984).

29. J.-F. Terral, A. Durand, C. Newton, S. Ivorra, Archéo-biologie de la domestication de l’olivier en Méditerranée occidentale : de la remise en cause d’une histoire dogmatique à la révélation de son irrigation médiévale. Etudes, 1–19 (2009).

30. G. Besnard, A. E. Bakkali, H. Haouane, D. Baali-Cherif, A. Moukhli, B. Khadari, Population genetics of Mediterranean and Saharan olives: Geographic patterns of differentiation and evidence for early generations of admixture. Ann. Bot. 112, 1293–1302 (2013).

31. B. Khadari, A. El Bakkali, Primary Selection and Secondary Diversification: Two Key Processes in the History of Olive Domestication. International Journal of Agronomy 2018, 1–9 (2018).

32. G. Besnard, J. F. Terral, A. Cornille, On the origins and domestication of the olive: A review and perspectives. Ann. Bot. 121, 385–403 (2018).

33. M. Gros-Balthazard, G. Besnard, G. Sarah, Y. Holtz, J. Leclercq, S. Santoni, D. Wegmann, S. Glémin, B. Khadari, Evolutionary transcriptomics reveals the origins of olives and the genomic changes associated with their domestication. Plant J. 100, 143–157 (2019).

34. L. Zunino, P. Cubry, G. Sarah, P. Mournet, A. El Bakkali, L. Aqbouch, S. Sidibé-Bocs, E. Costes, B. Khadari, Genomic evidence of genuine wild versus admixed olive populations evolving in the same natural environments in western Mediterranean Basin. PLoS One 19, e0295043 (2024).

35. D. Zohary, P. Spiegel-roy, Beginnings of Fruit Growing in the Old World. American Association for the Advancement of Science 187, 319–327 (1975).

36. R. Rubio de Casas, G. Besnard, P. Schönswetter, L. Balaguer, P. Vargas, Extensive gene flow blurs phylogeographic but not phylogenetic signal in Olea europaea L. Theor. Appl. Genet. 113, 575–583 (2006).

37. C. M. Diez, I. Trujillo, N. Martinez-Urdiroz, D. Barranco, L. Rallo, P. Marfil, B. S. Gaut, Olive domestication and diversification in the Mediterranean Basin. New Phytol. 206, 436–447 (2015).

38. D. N. Karger, O. Conrad, J. Böhner, T. Kawohl, H. Kreft, R. W. Soria-Auza, N. E. Zimmermann, H. P. Linder, M. Kessler, Climatologies at high resolution for the earth’s land surface areas. Scientific Data 4, 1–20 (2017).

39. P. Brun, N. Zimmermann, C. Hari, L. Pellissier, D. Karger, Global climate-related predictors at kilometer resolution for the past and future. *Earth Syst*. Sci. Data 14, 5573–5603 (2022).

40. I. Julca, M. Marcet-Houben, F. Cruz, J. Gómez-Garrido, B. S. Gaut, C. M. Díez, I. G. Gut, T. S. Alioto, P. Vargas, T. Gabaldón, Genomic evidence for recurrent genetic admixture during the domestication of Mediterranean olive trees (Olea europaea L.). BMC Biol. 18, 1–25 (2020).

41. P. Danecek, A. Auton, G. Abecasis, C. A. Albers, E. Banks, M. A. DePristo, R. E. Handsaker, G. Lunter, G. T. Marth, S. T. Sherry, G. McVean, R. Durbin, 1000 Genomes Project Analysis Group, The variant call format and VCFtools. Bioinformatics 27, 2156–2158 (2011).

42. E. Frichot, O. François, LEA: An R package for landscape and ecological association studies. Methods Ecol. Evol. 6, 925–929 (2015).

43. K. E. Lotterhos, M. C. Whitlock, The relative power of genome scans to detect local adaptation depends on sampling design and statistical method. Mol. Ecol. 24, 1031–1046 (2015).

44. S. Purcell, C. Chang, PLINK (2015; www.cog-genomics.org/plink/2.0/).

45. C. C. Chang, C. C. Chow, L. C. Tellier, S. Vattikuti, S. M. Purcell, J. J. Lee, Second-generation PLINK: rising to the challenge of larger and richer datasets. Gigascience 4, 7 (2015).

46. F. Privé, K. Luu, B. J. Vilhjálmsson, M. G. B. Blum, Performing Highly Efficient Genome Scans for Local Adaptation with R Package pcadapt Version 4. Mol. Biol. Evol. 37, 2153–2154 (2020).

47. B. Dauphin, EnvironmentalData, WSL/SLF GitLab (2024). https://gitlabext.wsl.ch/dauphin/EnvironmentalData.

48. K. Caye, B. Jumentier, J. Lepeule, O. François, LFMM 2: Fast and accurate inference of gene-environment associations in genome-wide studies. Mol. Biol. Evol. 36, 852–860 (2019).

49. J. D. Storey, A. J. Bass, A. Dabney, D. Robinson, G. Warnes, Qvalue: Q-Value Estimation for False Discovery Rate Control (2025; https://bioconductor.org/packages/qvalue.).

50. A. R. Quinlan, I. M. Hall, BEDTools: a flexible suite of utilities for comparing genomic features. Bioinformatics 26, 841–842 (2010).

51. M. Carlson, GO.Db: A Set of Annotation Maps Describing the Entire Gene Ontology (2022).

52. H. Pagès, M. Carlson, AnnotationDbi: Manipulation of SQLite-Based Annotations in Bioconductor (2023; https://bioconductor.org/packages/AnnotationDbi).

53. T. Wu, E. Hu, S. Xu, M. Chen, P. Guo, Z. Dai, T. Feng, L. Zhou, W. Tang, L. Zhan, X. Fu, S. Liu, X. Bo, G. Yu, clusterProfiler 4.0: A universal enrichment tool for interpreting omics data. Innovation (Camb) 2, 100141 (2021).

54. G. Yu, Enrichplot: Visualization of Functional Enrichment Result (2023; https://yulab-smu.top/biomedical-knowledge-mining-book/).

55. P. Cingolani, A. Platts, L. L. Wang, M. Coon, T. Nguyen, L. Wang, S. J. Land, X. Lu, D. M. Ruden, A program for annotating and predicting the effects of single nucleotide polymorphisms, SnpEff: SNPs in the genome of Drosophila melanogaster strain w1118; iso-2; iso-3. Fly 6, 80–92 (2012).

56. J. Gower, Generalized procrustes analysis. Psychometrika 40, 33–51 (1975).

57. J. Oksanen, G. L. Simpson, F. G. Blanchet, R. Kindt, P. Legendre, P. R. Minchin, R. B. O’Hara, P. Solymos, M. H. H. Stevens, E. Szoecs, H. Wagner, M. Barbour, M. Bedward, B. Bolker, D. Borcard, G. Carvalho, M. Chirico, M. De Caceres, S. Durand, H. B. A. Evangelista, R. FitzJohn, M. Friendly, B. Furneaux, G. Hannigan, M. O. Hill, L. Lahti, D. McGlinn, M.-H. Ouellette, E. Ribeiro Cunha, T. Smith, A. Stier, C. J. F. Ter Braak, J. Weedon, Vegan: Community Ecology Package (2022; https://CRAN.R-project.org/package=vegan).

58. D. Grivet, F. Sebastiani, R. Alía, T. Bataillon, S. Torre, M. Zabal-Aguirre, G. G. Vendramin, S. C. González-Martínez, Molecular footprints of local adaptation in two Mediterranean conifers. Mol. Biol. Evol. 28, 101–116 (2011).

59. J. A. Ramírez-Valiente, L. Santos del Blanco, R. Alía, J. J. Robledo-Arnuncio, J. Climent, Adaptation of Mediterranean forest species to climate: Lessons from common garden experiments. J. Ecol. 110, 1022–1042 (2022).

60. P. Fiorino, S. Mancuso, Differential thermal analysis, supercooling and cell viability in organs of Olea europaea at subzero temperatures. Adv. Hortic. Sci. 14, 23–27 (2000).

61. R. Petruccelli, G. Bartolini, T. Ganino, S. Zelasco, L. Lombardo, E. Perri, M. Durante, R. Bernardi, Cold Stress, Freezing Adaptation, Varietal Susceptibility of Olea europaea L.: A Review. Plants 11 (2022).

62. W. Köppen, Versuch einer Klassifikation der Klimate, vorzugsweise nach ihren Beziehungen zur Pflanzenwelt. Geogr. Z. 6, 593–611 (1900).

63. P. Lionello, F. Abrantes, L. Congedi, F. Dulac, M. Gacic, D. Gomis, C. Goodess, H. Hoff, H. Kutiel, J. Luterbacher, S. Planton, M. Reale, K. Schröder, M. Vittoria Struglia, A. Toreti, M. Tsimplis, U. Ulbrich, E. Xoplaki, “Introduction: Mediterranean Climate—Background Information” in The Climate of the Mediterranean Region, P. Lionello, Ed. (Elsevier, Oxford, 2012), pp. xxxv–xc.

64. M. Podgornik, J. Fantinič, T. Pogačar, V. Zupanc, Analysis of Olive tree flowering behavior based on thermal requirements: A case study from the northern Mediterranean region. Climate 13, 156 (2025).

65. K. Fujita, H. Inui, Review: Biological functions of major latex-like proteins in plants. Plant Sci. 306, 110856 (2021).

66. I. Ventura, L. Brunello, S. Iacopino, M. C. Valeri, G. Novi, T. Dornbusch, P. Perata, E. Loreti, Arabidopsis phenotyping reveals the importance of alcohol dehydrogenase and pyruvate decarboxylase for aerobic plant growth. Sci. Rep. 10, 16669 (2020).

67. J. Rojo, F. Orlandi, A. Ben Dhiab, B. Lara, A. Picornell, J. Oteros, M. Msallem, M. Fornaciari, R. Pérez-Badia, Estimation of chilling and heat accumulation periods based on the timing of Olive pollination. Forests, doi: 10.3390/f11080835 (2020).

68. L.-F. Li, S. A. Cushman, Y.-X. He, Y. Li, Genome sequencing and population genomics modeling provide insights into the local adaptation of weeping forsythia. Horticulture Research 7, 1–12 (2020).

69. O. M. Nedukha, Callose: Localization, functions, and synthesis in plant cells. Cytol. Genet. 49, 49–57 (2015).

70. G. Miller, N. Suzuki, S. Ciftci-Yilmaz, R. Mittler, Reactive oxygen species homeostasis and signalling during drought and salinity stresses. Plant Cell Environ. 33, 453–467 (2010).

71. X. Liu, T. Gao, C. Liu, K. Mao, X. Gong, C. Li, F. Ma, Fruit crops combating drought: Physiological responses and regulatory pathways. Plant Physiol. 192, 1768–1784 (2023).

72. R. D. H. Barrett, D. Schluter, Adaptation from standing genetic variation. Trends Ecol. Evol. 23, 38–44 (2008).

73. A. Kremer, O. Ronce, J. J. Robledo-Arnuncio, F. Guillaume, G. Bohrer, R. Nathan, J. R. Bridle, R. Gomulkiewicz, E. K. Klein, K. Ritland, A. Kuparinen, S. Gerber, S. Schueler, Long-distance gene flow and adaptation of forest trees to rapid climate change. Ecol. Lett. 15, 378–392 (2012).

74. R. J. Petit, C. Bodénès, A. Ducousso, G. Roussel, A. Kremer, Hybridization as a mechanism of invasion in oaks. New Phytol. 161, 151–164 (2004).

75. R. J. Petit, A. Hampe, Some evolutionary consequences of being a tree. Annu. Rev. Ecol. Evol. Syst. 37, 187–214 (2006).

76. J. Kulmuni, B. Wiley, S. Otto, On the fast track: hybrids adapt more rapidly than parental populations in a novel environment. Evol. Lett. 8, 128–136 (2023).

77. S. Lavee, D. Zohary, The potential of genetic diversity and the effect of geographically isolated resources in olive breeding. Isr. J. Plant Sci. 59, 3–13 (2011).

78. Y. Chen, Z. Jiang, P. Fan, P. G. P. Ericson, G. Song, X. Luo, F. Lei, Y. Qu, The combination of genomic offset and niche modelling provides insights into climate change-driven vulnerability. Nat. Commun. 13, 4821 (2022).

