## Supplementary Information for "Genomic insights into the local adaptation of spontaneously occurring populations of olive trees (*Olea europaea ssp. europaea var. sylvestris*) in the Mediterranean Basin"

**2 AGAP Institut, Univ Montpellier, CIRAD, INRAE, Institut Agro, Montpellier, France**

**3 DIADE, Univ Montpellier, IRD, CIRAD, Montpellier, France**

**4 CIRAD, UMR AGAP Institut, F-34398 Montpellier, France**

**5 INRA, UR Amélioration des Plantes et Conservation des Ressources Phytogénétiques, Meknès, Morocco**

**6 Conservatoire Botanique National Méditerranéen (CBNMed), UMR AGAP Institut, Montpellier, France**

**Abstract**:

Living organisms are increasingly threatened by significant environmental changes, primarily driven by human-induced global alterations. Understanding and forecasting species adaptive responses to these environmental changes is therefore crucial for enhancing conservation efforts. In the Mediterranean Basin (MB), the average temperature is rising at a rate 20% faster than the global average, placing regional biodiversity at heightened risk. The study of local adaptation is thus particularly relevant. In the western MB, spontaneously occurring olive populations consist of both wild olives and admixed resulting of crop-to-wild gene flow. The wild is likely harbouring adaptive genetic variation shaped by local environmental pressures. In this study, we analysed target genomic sequencing data from spontaneously occurring populations (comprising both wild and admixed trees) as well as cultivated olive trees. We performed selective sweep and GEA analyses in wild olives to identify genomic regions associated with environmental variables. Our findings identified signatures of adaptation particularly associated with precipitation and temperature. Notably, admixed individuals retained many wild candidate SNPs in their genomes suggesting that they might retain a certain level of adaptation to their local environment. Those results are in line with recent studies suggesting hybrids could adapt more rapidly to novel environments than their parental populations.

**The following Supporting Information is available for this article:**

### FIGURE

**Figure S1. Geographical distribution of sampling of the spontaneously occurring western populations of Mediterranean Basin of *O. europaea* subsp. *europaea* var *sylvestris* for this study.**

**Figure S2**. **Principal Component Analysis of climatic data of 26 naturally occurring populations of olive trees in western Mediterranean Basin.**

**Figure S3. Result of the population structure analyses performed on the genome-wide SNPs diversity of spontaneously occurring populations olives (n=359) and cultivated olives (n=145) from western Mediterranean Basin.**

**Figure S4**. **Proportion of individuals characterized as wild or admixed olive genotypes in the spontaneously occurring western populations of Mediterranean Basin.**

**Figure S5. Genome Scan for Selection: PCA resulting from pcadapt analysis in wild olive trees of western MB.**

**Figure S6. Genome scan for selection in western MB wild olive trees using pcadapt: Quantile–quantile (Q–Q) plot of observed vs. expected −log₁₀(p-values).**

**Figure S7. Genome scan for selection in western MB wild olive trees using pcadapt: histogram of p-value distribution across all SNPs.**

**Figure S8.** **Principal Component analysis of all environmental data for the 26 sites sampled**

**Figure S9.** **Pearson correlation matrix of all environmental data per site.**

**Figure S10.** **Principal Component analysis of reduce correlations environmental data (<75% Pearson Correlation) for the 26 sites sampled**

**Figure S11. Manhattan plots on chromosome of genomic–environment associations (GEAs) for wild olive trees from the western Mediterranean Basin.**

**Figure S12. Genome scan for local adaptation in western MB wild olive trees using LFMM: Quantile–quantile (Q–Q) plots for each environmental variable tested in the genomic–environment association analysis.**

**Figure S13. Genome scan for local adaptation in western MB wild olive trees using LFMM: Histograms of p-value distributions for each environmental variable tested in the genomic–environment association analysis.**

**Figure S14. Gene ontology (GO) term enrichment analysis based on SNPs associated with environmental variables detected by LFMM for western MB wild olive trees.**

### TABLE

**Table S1.** **Summary information of olive tree dataset for this study.**

**Table S2.** **List of baits used for this study**.

**Table S3.** **Summary information of all environmental data of 26 naturally occurring populations of *O. europaea* used in this study.**

**Table S4. Result of the assignment of spontaneously occurring populations olives from Western MB to genetic clusters wild or admixed.**

**Table S5**. **Summary of outliers SNPs detected by selective sweep detection using pcadapt for wild olives *O. europaea* L.**

**Table S6. Q-value threshold retained per variable tested during genome-environment association analysis for wild genotypes.**

**Table S7. Summary of outliers SNPs detected by GEA analysis using LFMM for wild olives *O. europaea* L.**

**Table S8. Outliers SNPs resulting from GEA analysis retained for Procrustes analysis.**

**Table S9. GEA and Selective Sweep Outliers with High SNPeff Impact**

**Table S10. BLAST Results for outliers SNPs detected by GEA for Bio12 and Bio19.**

**Method S1. Selection of sampling sites according to climatic data clusters.**

**Method S1. Selection of sampling sites according to climatic data clusters.** To prevent redundancy in climatic data per population, for each potential site pre-selected according to the criteria given in Zunino et al (2024) we extracted climatic data from WorldClim version 2.1 (Fick & Hijmans 2017; Harris *et al.* 2020). For each site, we examined the following climatic variables from 1971 to 2000: the average monthly minimum temperature of the coldest quarter, calculated with the monthly data, and the average monthly precipitation of the driest quarter (Bio17). These variables were chosen according to the biology of the olive tree, with the chilling requirement parameters and the precipitation requirements of olive trees. (Haberman *et al.* 2017; Alfieri *et al.* 2019). Then, the sites were grouped in six clusters based on these data using a kmeans clustering approach, a machine learning method whose aim is to regroup similar data in clusters, implemented with the kmeans function from R version 4.2.0, (R Core Team 2022). Within each cluster, we selected one to two potential sites to avoid redundancy of climatic data between geographically closed populations. After sampling all populations, we extracted the 19 Bioclimatic variables of each site from the CHELSA database, which contains recent climatic variables from 1981 to 2010 (Karger *et al.* 2017; Brun *et al.* 2022). A final assessment of climatic data heterogeneity at the sites was conducted using a Principal Component Analysis (PCA; Fig. S2).

### FIGURE

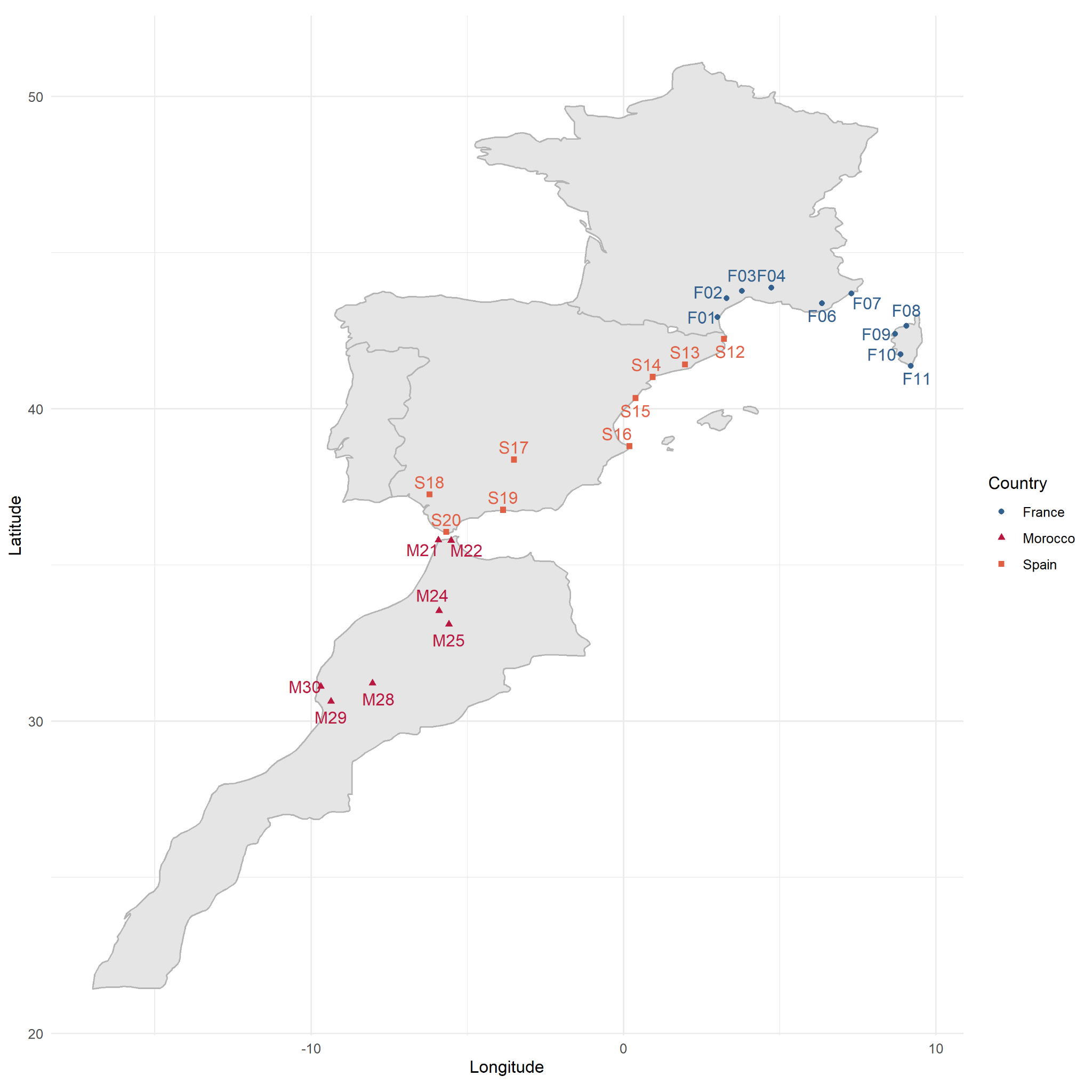

**Figure S1**. **Geographical distribution of sampling of the spontaneously occurring western populations of Mediterranean Basin of *O. europaea* subsp. *europaea* var *sylvestris* for this study.** The native populations of olives tree *O. europaea* are found in France (blue circle), in Spain (orange square) and in Morocco (red triangle).

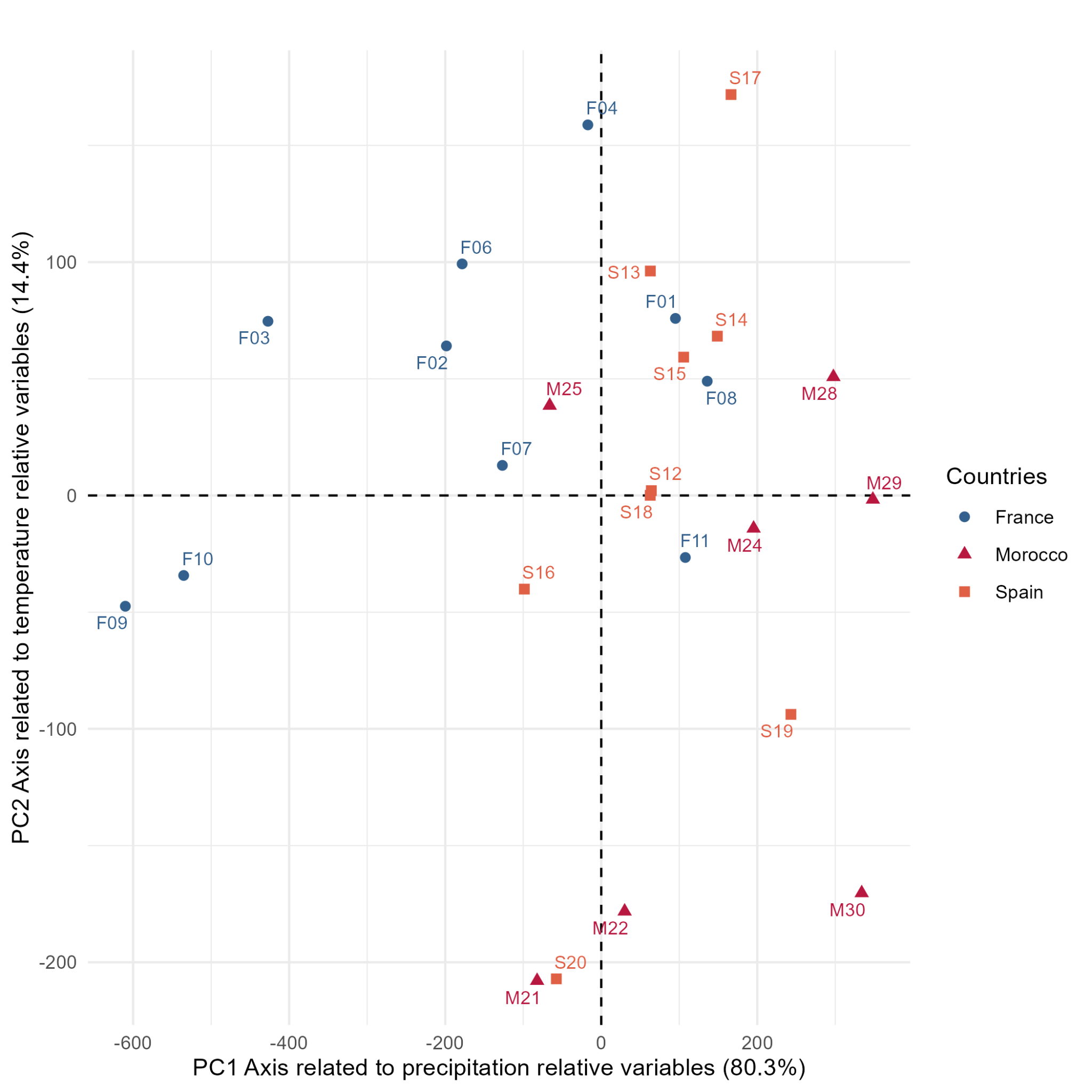

**Figure S2**. **Principal Component Analysis of climatic data of 26 naturally occurring populations of olive trees in western Mediterranean Basin.** The native populations of olives tree *O. europaea* are found in France (blue circle), in Spain (orange square) and in Morocco (red triangle). The first dimension is correlated to the precipitation-related variables and the second dimension to the temperature-related variables, from 1981 to 2010 in CHELSA V2 (Karger *et al.* 2017).

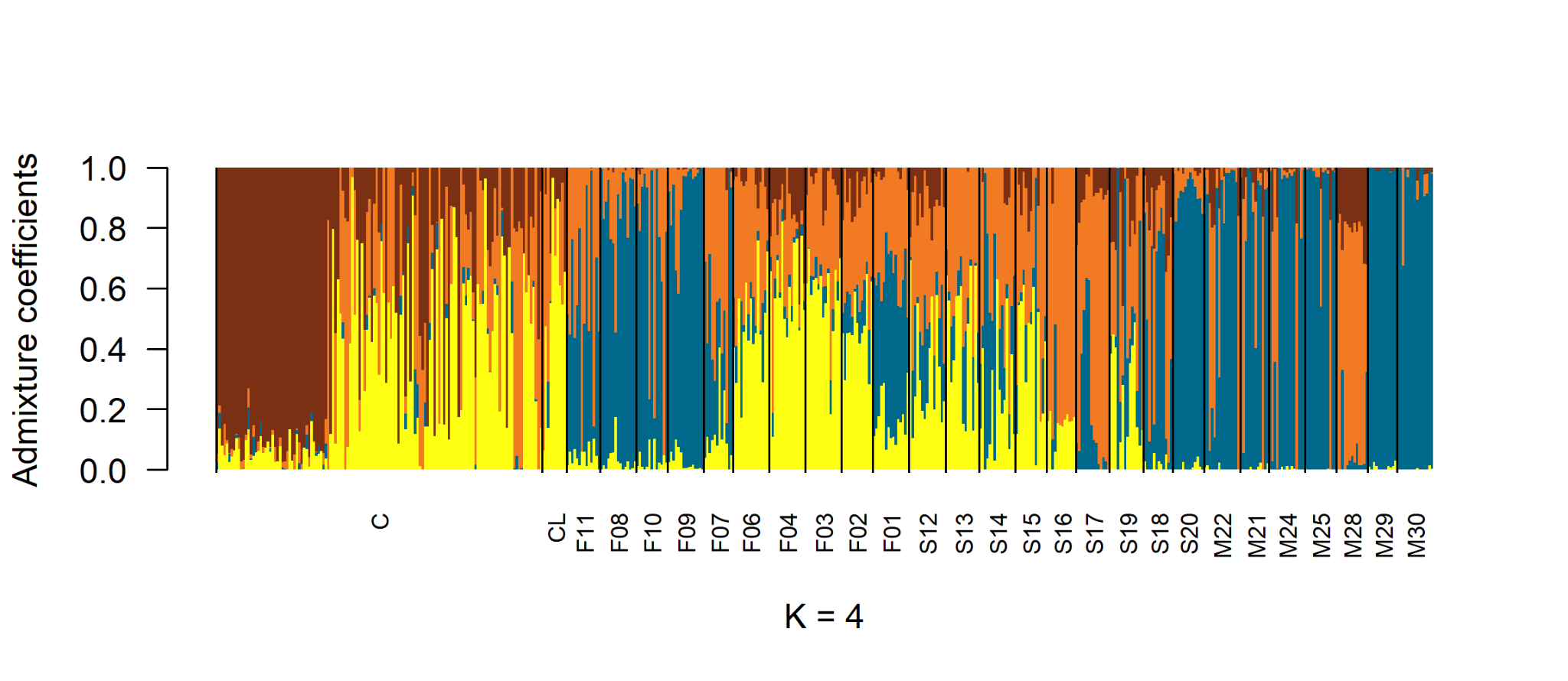

**Figure S3. Result of the population structure analyses performed on the genome-wide SNPs diversity of spontaneously occurring populations olives (n=359) and cultivated olives (n=145) from western Mediterranean Basin.** Genetic structure inferred by sNMF, each horizontal bar indicates individual assignment to a genetic cluster with K being the number of genetic clusters, here K=4. C referred to cultivated from the Worldwide Olive Germplasm Bank of Marrakech (WOGBM) and CL referred to cultivated from France collection in Porquerolles and in the Technical Center of Olives (CTO)

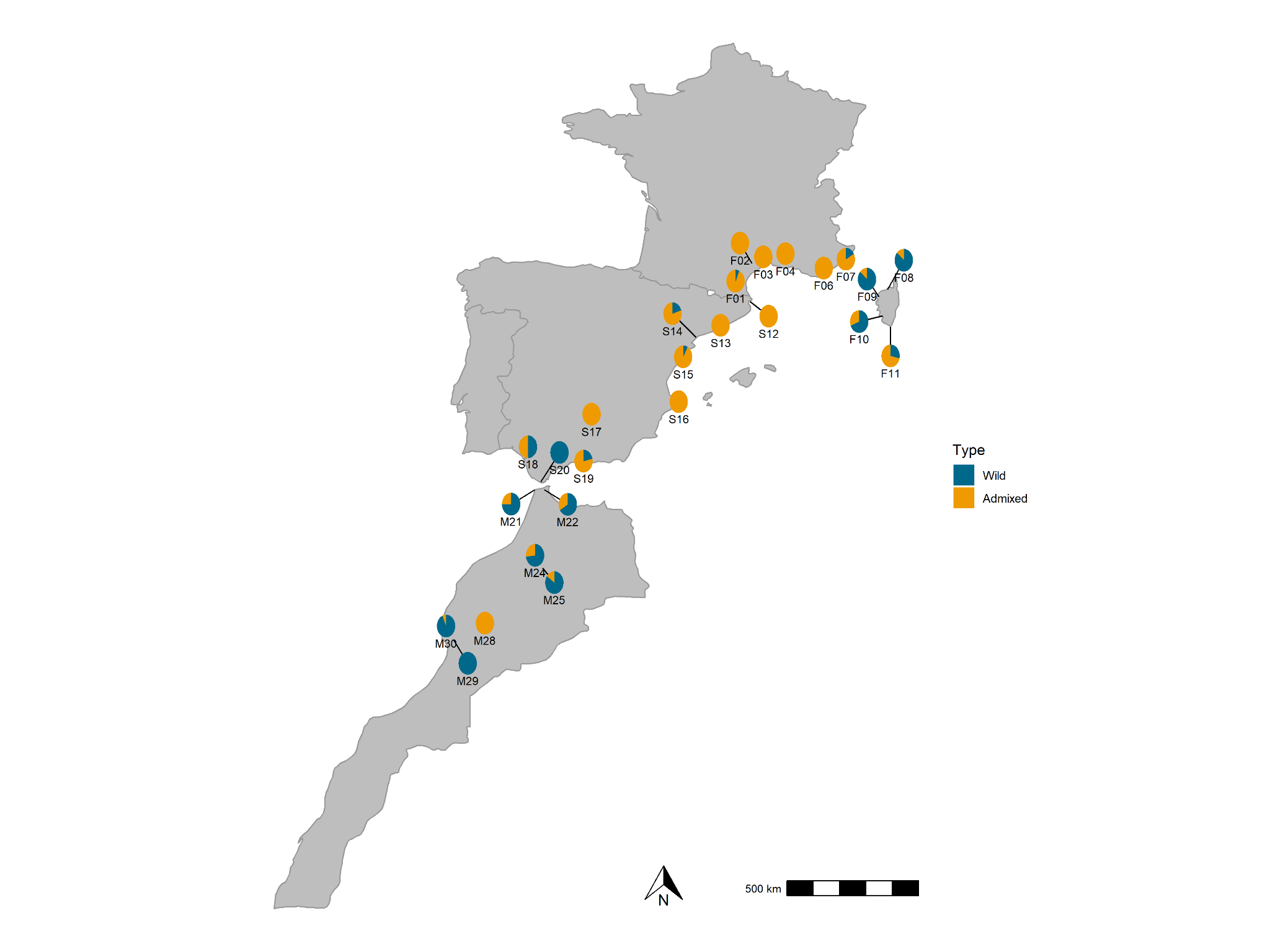

**Figure S4**. **Proportion of individuals characterized as wild or admixed olive genotypes in the spontaneously occurring western populations of Mediterranean Basin.** The pie-charts report the proportions of wild individuals (in blue) and admixed individuals (in yellow).

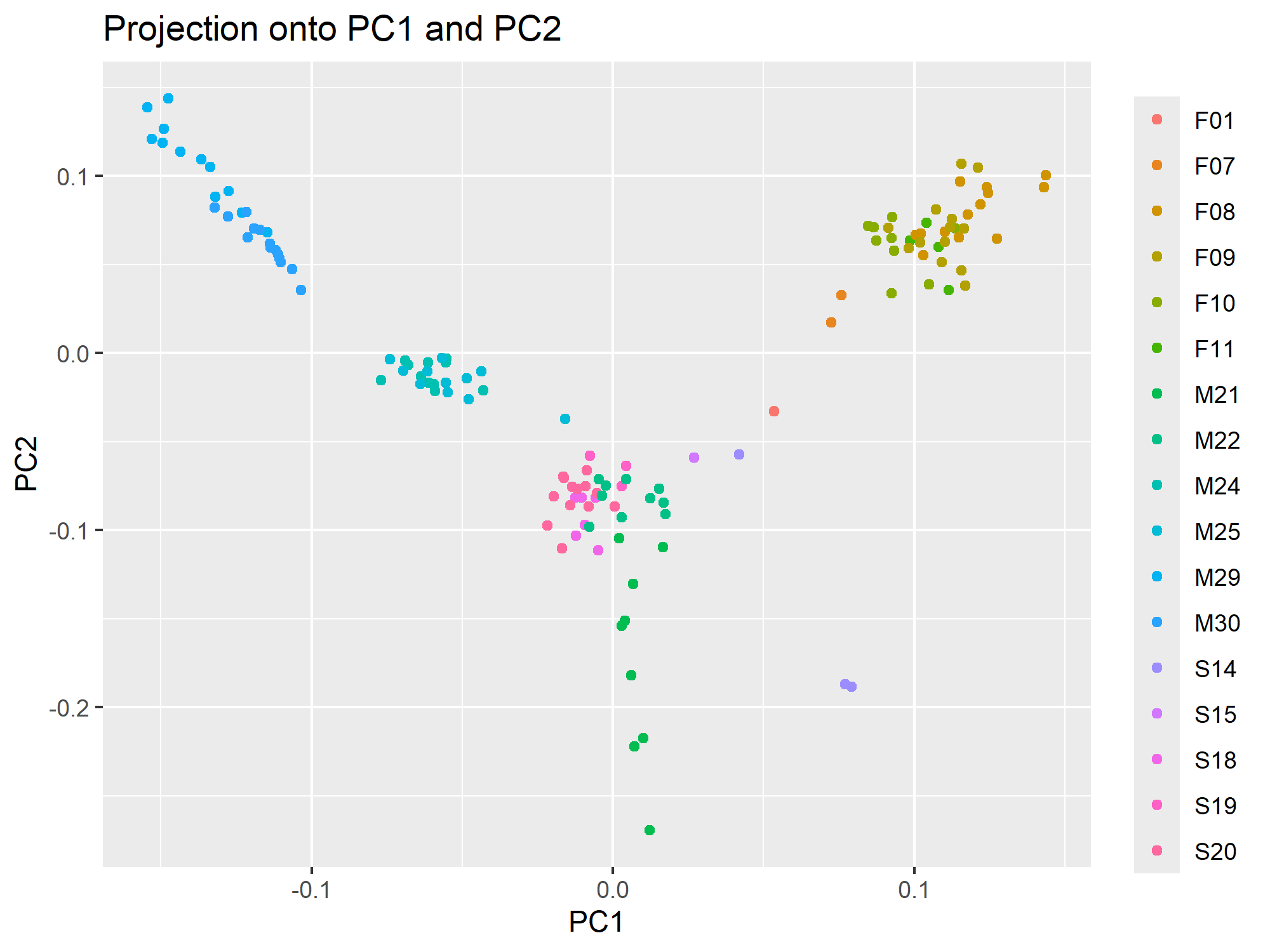

**Figure S5. Genome Scan for Selection: PCA resulting from pcadapt analysis in wild olive trees of western MB.** Each dot represents a SNP, and each color corresponds to a sampling site.

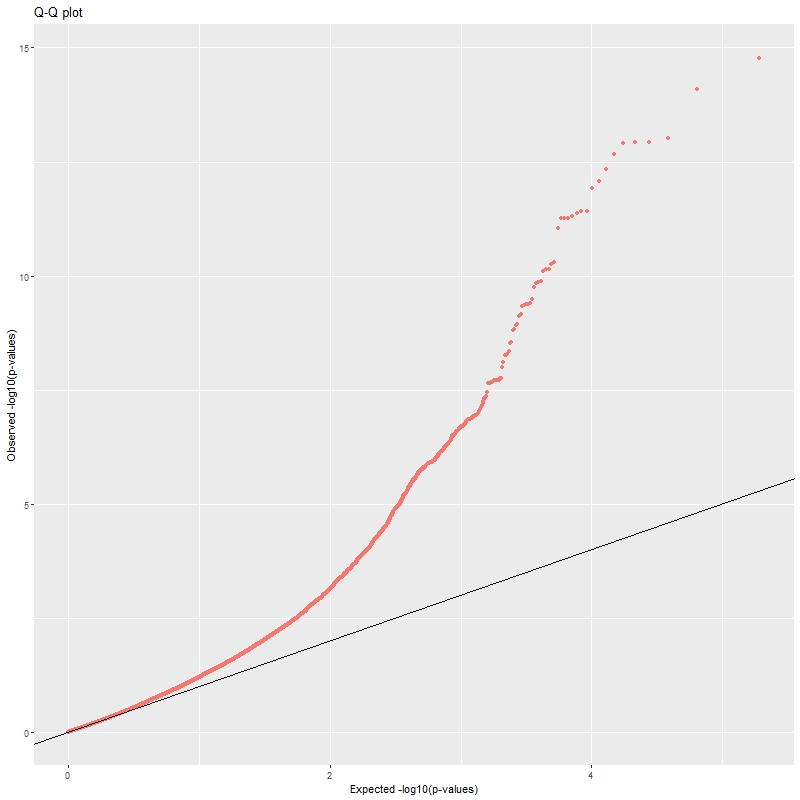

**Figure S6. Genome scan for selection in western MB wild olive trees using pcadapt: Quantile–quantile (Q–Q) plot of observed vs. expected −log₁₀(p-values).**

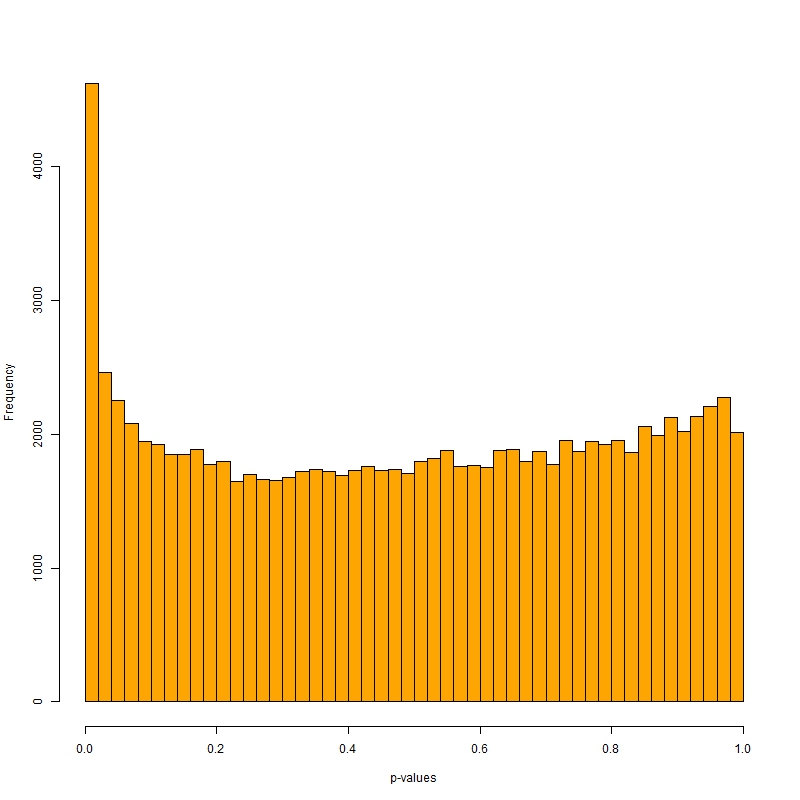

**Figure S7. Genome scan for selection in western MB wild olive trees using pcadapt: histogram of p-value distribution across all SNPs.**

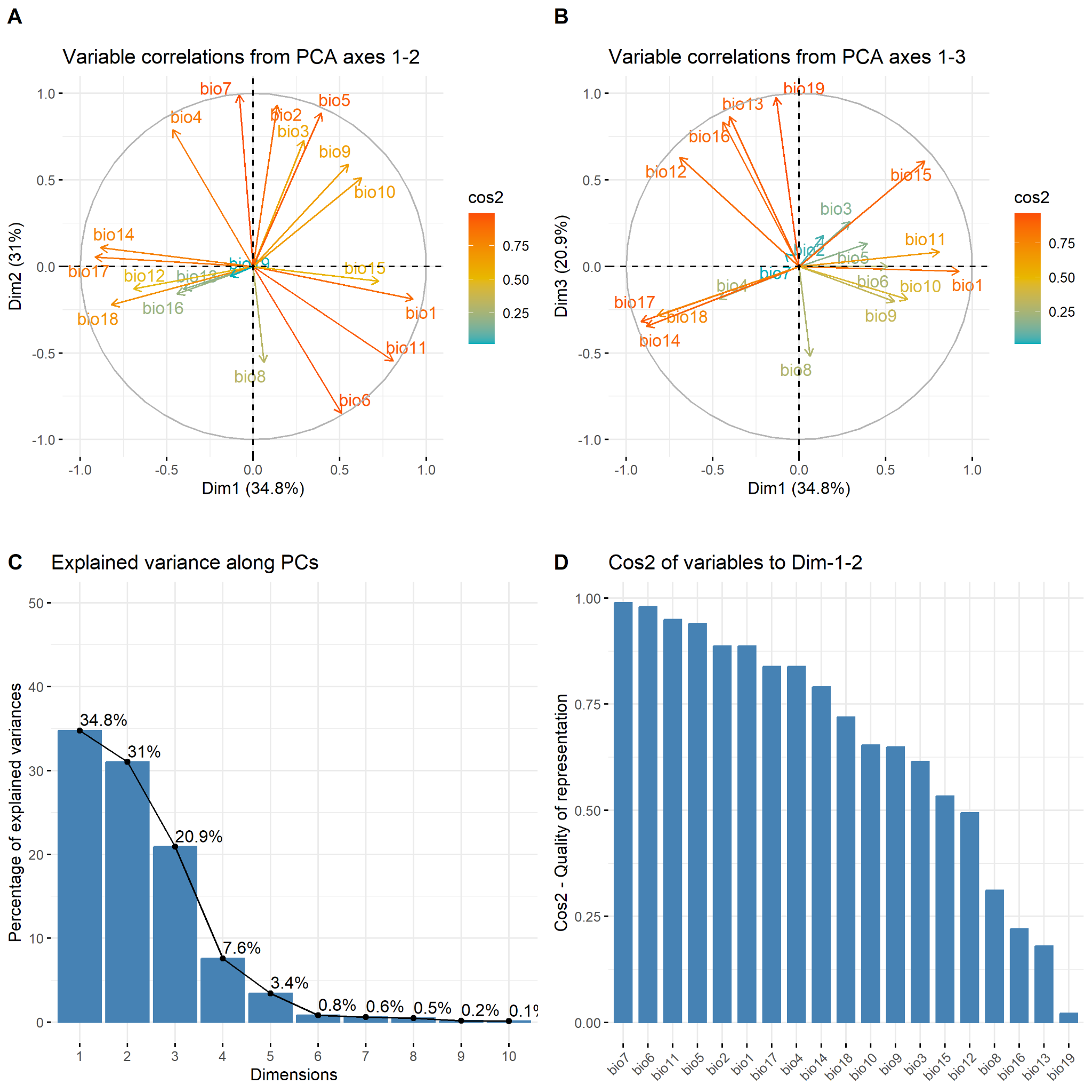

**Figure S8.** **Principal Component analysis of all environmental data for the 26 sites sampled** with (A) the variable correlations axes 1-2 (B) the variable correlations axes 1-3 (C) the explained variance for each dimension (D) the Cos2 of variables for Dim-1-2. Analysis based on Benjamin Dauphin script available here: <https://github.com/bcm-uga/SSMPG2022>

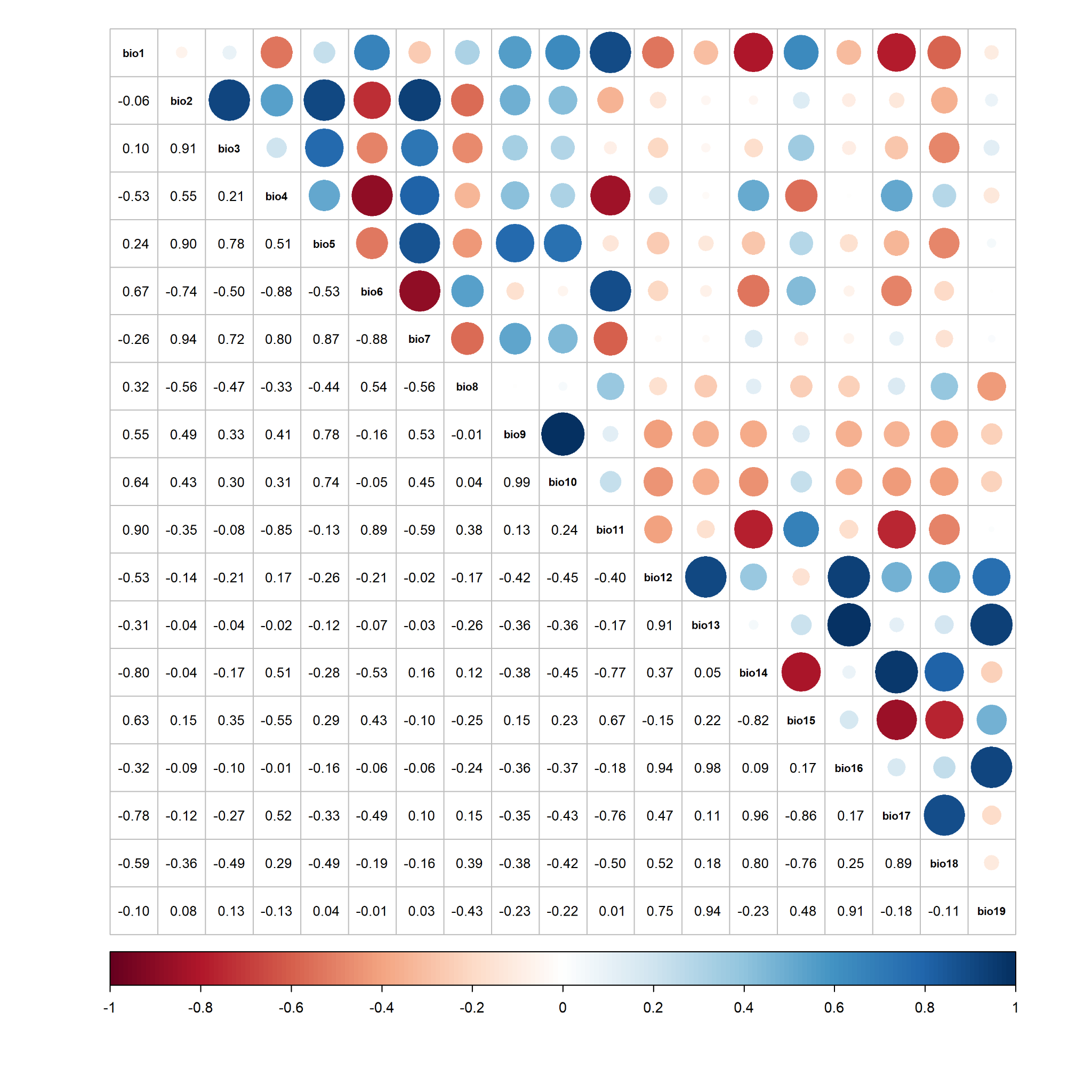

**Figure S9.** **Pearson correlation matrix of all environmental data per site.**

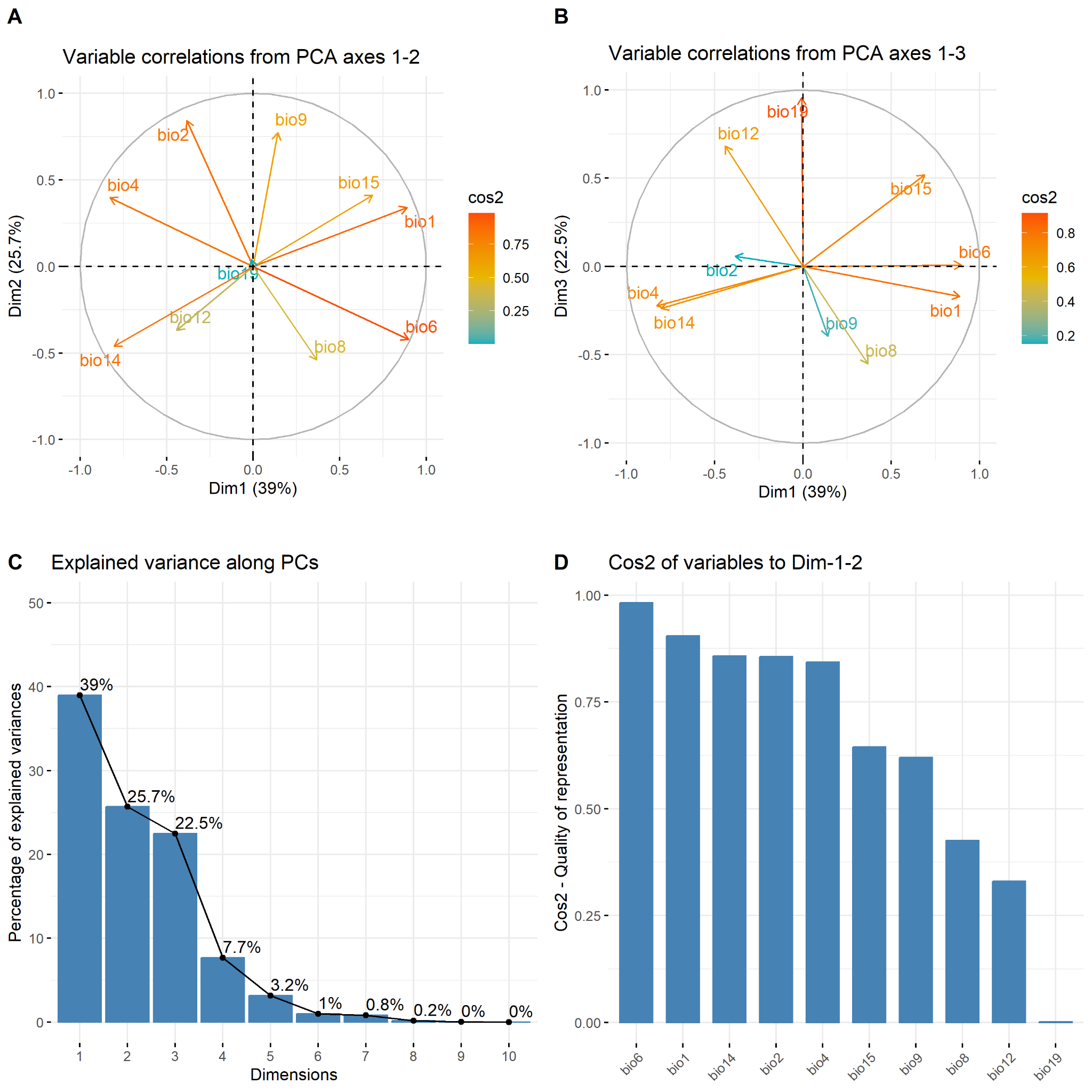

**Figure S10.** **Principal Component analysis of reduce correlations environmental data (<75% Pearson Correlation) for the 26 sites sampled** with (A) the variables correlation axes 1-2 (B) the variable correlations axes 1-3 (C) the explained variance for each dimension (D) the Cos2 of variables for Dim-1-2.

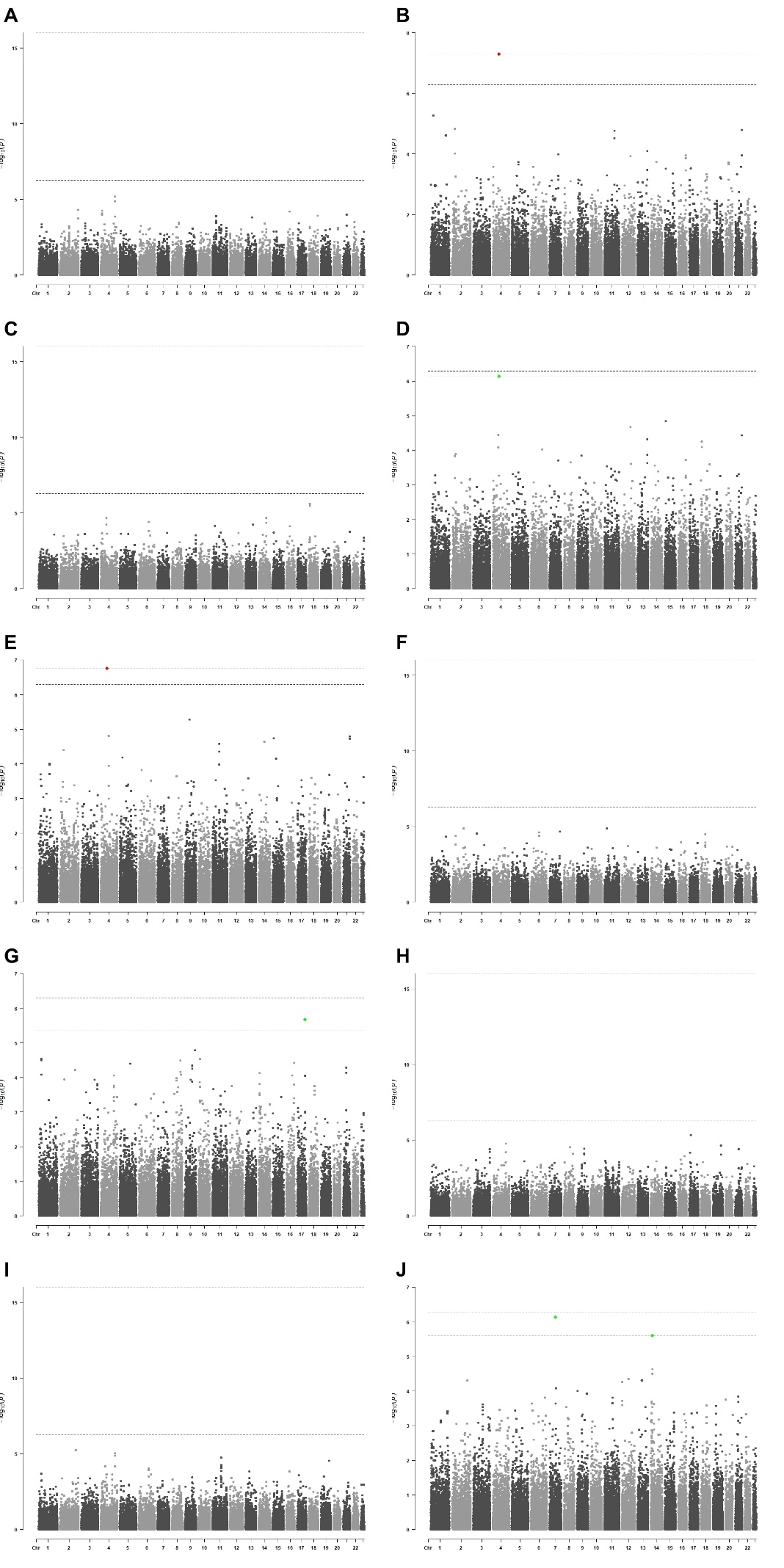

**Figure S11. Manhattan plots on chromosome of genomic–environment associations (GEAs) for wild olive trees from the western Mediterranean Basin.** Each point represents a SNP. Yellow points indicate minor outlier SNPs with q-value < 0.1. Pink points indicate major outlier SNPs with p-value < 0.05 after Bonferroni correction. Environmental variables are shown in the following order: A) “bio1”, B) “bio2”, C) “bio4”, D) “bio6”, E) “bio8”, F) “bio9”, G) “bio12”, H) “bio14”, I) “bio15”, and “J) bio19”.

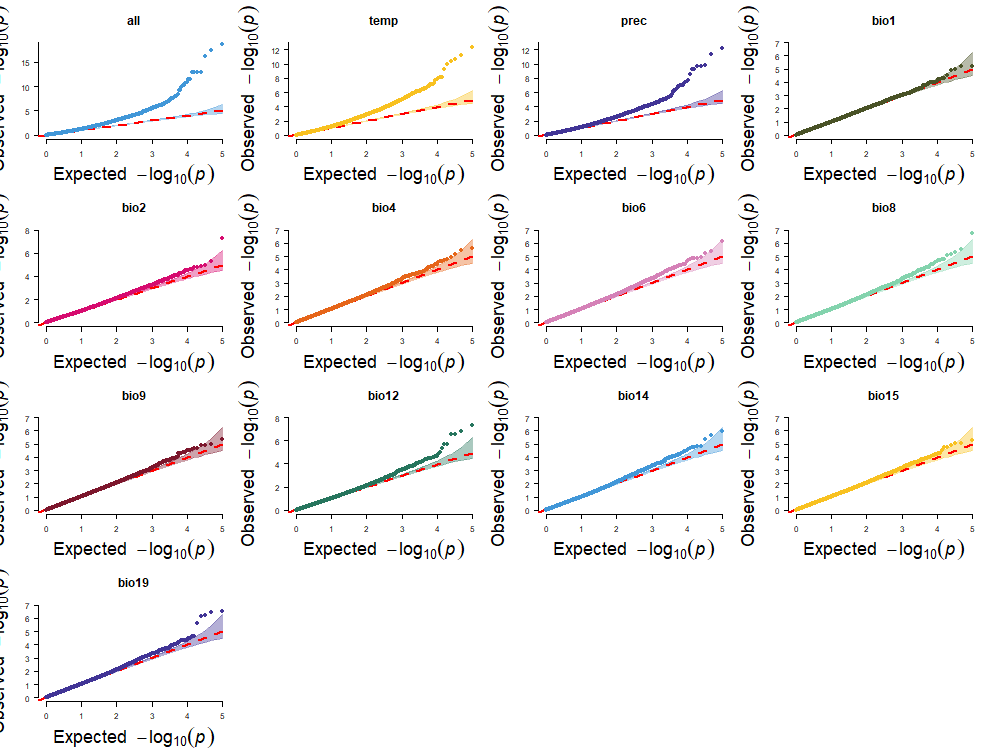

**Figure S12. Genome scan for local adaptation in western MB wild olive trees using LFMM: Quantile–quantile (Q–Q) plots for each environmental variable tested in the genomic–environment association analysis.** Variables shown in order: “all”, “temperature”, “precipitation”, “bio1”, “bio2”, “bio4”, “bio6”, “bio8”, “bio9”, “bio12”, “bio14”, “bio15”, and “bio19”.

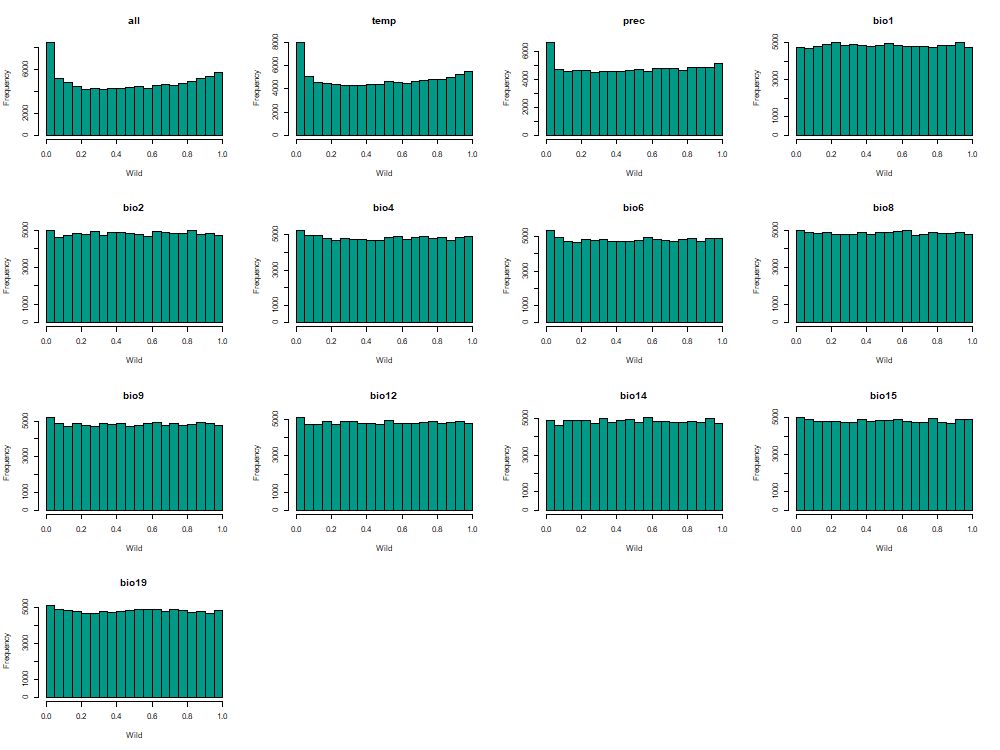

**Figure S13. Genome scan for local adaptation in western MB wild olive trees using LFMM: Histograms of p-value distributions for each environmental variable tested in the genomic–environment association analysis.** Variables are shown in the following order: “all”, “temperature”, “precipitation”, “bio1”, “bio2”, “bio4”, “bio6”, “bio8”, “bio9”, “bio12”, “bio14”, “bio15”, and “bio19”.

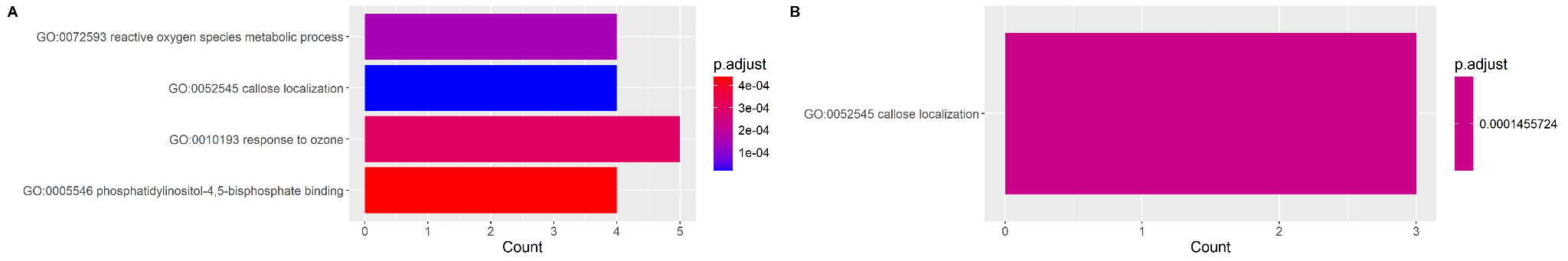

**Figure S14. Gene ontology (GO) term enrichment analysis based on SNPs associated with environmental variables detected by LFMM for western MB wild olive trees.** (A) SNPs associated to “All” climate variable (B) SNPs associated to “Temperature”related variable.

### TABLE

**Table S1**. **Summary information of olive tree dataset for this study.**

MB - Mediterranean Basin, WOGBM - World Olive Germplasm of Bank of Marrakech POR: Porquerolles collection, CTO: Technical Center of Olive tree

**Table S2.** **List of baits used for this study (chromosome and scaffolds)**. The baits were designed based on annotated genes of *Olea europaea* var. *europaea* (cv. Farga) Oe9 genome assembly (Julca *et al.* 2020).

**Table S3.** **Summary information of all environmental data of 26 naturally occurring populations of *O. europaea* used in this study.** Environmental data are from 1981 to 2010, extracted from CHELSA V2 database (Karger *et al.* 2017)

*Bio1: Mean annual air temperature (°C), Bio2: Mean diurnal temperature arrange (°C), Bio3: Isothermality (°C), Bio4: Temperature seasonality (°C/100), Bio5:mean daily maximum air temperature of the warmest month (°C), Bio6: Mean daily minimum air temperature of the coldest month (°C), Bio7: annual range of air temperature (°C), Bio8: Mean daily air temperatures of the wettest quarter (°C), Bio9: Mean daily air temperatures of the driest quarter (°C), Bio10: mean daily mean air temperatures*

*of the warmest quarter (°C), Bio11: mean daily mean air temperatures of the coldest quarter (°C), Bio12: Annual precipitation amount (Kg.m-2.year-1), Bio13: precipitation amount of the wettest month (kg m-2, month-1), Bio14: Precipitation amount of the driest month (kg m-2 month-1), Bio15: Precipitation seasonality (kg.m-2), Bio16: mean monthly precipitation amount of the wettest quarter (kg m-2, month-1), Bio17: mean monthly precipitation amount of the driest quarter (kg m-2, month-1), Bio18: mean monthly precipitation amount of the warmest quarter (kg.m-2.month-1), Bio19: Mean monthly precipitation amount of the coldest quarter (kg.m-2.month-1).*

**Table S4. Result of the assignment of spontaneously occurring populations olives from Western MB to genetic clusters wild or admixed.** Individuals with an assignment probability ≥70% to C4 cluster were classified as genetically "wild", whereas individuals below this threshold (unassigned) or assigned to C1, C2 or C3 were categorized as "admixed”.

**Table S5**. **Summary of outliers SNPs detected by selective sweep detection using pcadapt for wild olives *O. europaea* L.**

The major outliers are detected using a *p-value* threshold < 0.05 altered with Bonferroni correction (alpha =0.05, n=96,648, adjusted threshold = 5.17e-07). A relaxed threshold of *q-value* < 0.1 allowed to identify additional minor outliers.

**Table S6. Q-value threshold retained per variable tested during genome-environment association analysis for wild genotypes and admixed genotypes.**

| Environmental variable | q-value |
| --- | --- |
| *all* | 1.03E-03 |
| *temperature* | 6.88E-04 |
| *precipitation* | 2.52E-04 |
| *Bio1* | NA |
| *Bio2* | 5.12E-08 |
| *Bio4* | NA |
| *Bio6* | 7.44E-07 |
| *Bio8* | 1.76E-07 |
| *Bio9* | NA |
| *Bio12* | 4.29E-06 |
| *Bio14* | NA |
| *Bio15* | NA |
| *Bio19* | 2.49E-06 |

**Table S7. Summary of outliers SNPs detected by GEA analysis using LFMM for wild olives *O. europaea* L.**

The major outliers are detected using a *p-value* threshold < 0.05 altered with Bonferroni correction (alpha =0.05, n=96,648, adjusted threshold = 5.17e-07). A relaxed threshold of *q-value* < 0.1 allowed to identify additional minor outliers. Bio1: Mean annual air temperature (°C), Bio2: Mean diurnal temperature arrange (°C), Bio4: Temperature seasonality (°C/100), Bio6: Mean daily minimum air temperature of the coldest month (°C), Bio8: Mean daily air temperatures of the wettest quarter (°C), Bio9: Mean daily air temperatures of the driest quarter (°C), Bio12: Annual precipitation amount (Kg.m-2.year-1), Bio14: Precipitation amount of the driest month (kg m-2 month-1), Bio15: Precipitation seasonality (kg.m-2), Bio19: Mean monthly precipitation amount of the coldest quarter (kg.m-2.month-1).

**Table S8. Outliers SNPs resulting from GEA analysis retained for Procrustes analysis.**

**Table S9. GEA and Selective Sweep Outliers with High SNPeff Impact**

**Table S10. BLAST Results for outliers SNPs detected by GEA for Bio12 and Bio19.**

References:
